# KS-WNK1 amplifies kidney tubule responsiveness to potassium via WNK body condensates

**DOI:** 10.1101/2021.03.12.435046

**Authors:** Cary R. Boyd-Shiwarski, Rebecca T. Beacham, Jared A. Lashway, Katherine E. Querry, Shawn E. Griffiths, Daniel J. Shiwarski, Sophia A. Knoell, Nga H. Nguyen, Lubika J. Nkashama, Melissa N. Valladares, Anagha Bandaru, Allison L. Marciszyn, Aylin R. Rodan, Chou-Long Huang, Sean D. Stocker, Ossama B. Kashlan, Arohan R. Subramanya

**Affiliations:** Department of Medicine, Renal-Electrolyte Division; Department of Medicine, Vascular Medicine Institute; Department of Cell Biology; Department of Computational and Systems Biology; Department of Neurobiology; Pittsburgh Center for Kidney Research University of Pittsburgh School of Medicine, Pittsburgh, PA, USA; Department of Bioengineering, Swanson School of Engineering, University of Pittsburgh, Pittsburgh PA, USA; Department of Internal Medicine, Division of Nephrology and Hypertension; Department of Human Genetics; Molecular Medicine Program University of Utah, Salt Lake City, UT, USA; Medical Service, VA Salt Lake City Healthcare System, Salt Lake City, UT, USA; Department of Internal Medicine, Division of Nephrology and Hypertension, University of Iowa Carver College of Medicine, Iowa City, IA, USA; VA Pittsburgh Healthcare System, Pittsburgh, PA, USA

## Abstract

To maintain potassium homeostasis, the kidney’s distal convoluted tubule (DCT) converts small changes in blood [K^+^] into robust effects on salt reabsorption. This process requires NaCl cotransporter (NCC) activation by WNK kinases. During hypokalemia, the Kidney-Specific WNK1 isoform (KS-WNK1) scaffolds the DCT-expressed WNK signaling pathway within biomolecular condensates of unknown function termed WNK bodies. Here, we show that KS-WNK1 amplifies the dynamic range of NCC activity in response to potassium imbalance, in part via WNK bodies. Targeted condensate disruption traps the WNK pathway, causing renal salt-wasting that is more pronounced in females. In humans, WNK bodies accumulate as plasma potassium falls below 4.0mmol/L, suggesting avid condensate-mediated salt reabsorption even when [K^+^] is low-normal. These data identify WNK bodies as signal amplifiers that mediate tubular potassium responsiveness, nephron sexual dimorphism, and blood pressure salt-sensitivity. Our results illustrate how condensate specialization can optimize a mammalian physiologic stress response that impacts human health.

## Introduction

During environmental stress, physiologic systems must sense imbalance and coordinate appropriately tuned responses that maintain homeostasis. An example of this is the distal nephron, a series of kidney tubule segments which sense and cooperatively maintain plasma potassium concentrations within the narrow physiologic window required for life^1^. During the stress of hypokalemia, a serine-threonine kinase cascade within the distal convoluted tubule (DCT) activates the thiazide-sensitive NaCl cotransporter (NCC; *SLC12A3*) via phosphorylation^2^. Hypokalemia-mediated NCC activation limits downstream delivery of sodium to the connecting tubule and collecting duct, diminishing distal voltage-dependent potassium secretion. This minimizes urinary cation losses to conserve total body potassium^3^. In contrast, hyperkalemia inhibits NCC, facilitating kaliuresis^4^.

With-No-Lysine (WNK) kinases are essential regulators of NCC phosphorylation and potassium homeostasis. During hypokalemia, the WNKs activate the kinases SPAK (*STK39*) and OSR1 (*OXSR1*), which phosphorylate NCC directly^1^. Conversely, hyperkalemia promotes NCC dephosphorylation^4,5^. The integration of these signals generates an inverse relationship between NCC phosphorylation status and plasma [K^+^]^6^. The importance of WNK signaling in potassium metabolism is evidenced by Familial Hyperkalemic Hypertension (FHHt), a Mendelian syndrome caused by overactivation of the DCT-expressed WNK signaling pathway, resulting in NCC hyperphosphorylation, salt-sensitive hypertension, and hyperkalemia that is cured with thiazide diuretics^7,8^.

The distal convoluted tubule features a unique complement of WNK-SPAK/OSR1 pathway gene products. WNK4 is the dominant DCT-expressed WNK kinase^9^. The full-length kinase-active “Long” isoform of WNK1 (L-WNK1) is also expressed in this nephron segment, though its abundance is low^10^. Instead, the most abundant WNK1 isoform in the DCT is a kidney-exclusive truncated gene product that lacks kinase activity. This “Kidney-Specific WNK1” (KS-WNK1) isoform requires an intragenic promoter located within intron 4 of the WNK1 gene (Fig 1A). This distal tubule-specific promoter drives the expression of an alternative first exon that replaces the L-WNK1 N-terminus and most of the kinase domain with 30 unique amino acids encoded by exon 4a^11^. Downstream from this sequence, L-WNK1 and KS-WNK1 are identical (Fig 1B). Exon 4a emerged during the evolutionary transition of vertebrates from water to land, a process that required robust kidney tubular transport to preserve electrolyte homeostasis^12,13^. Exon 4a is also critical for the formation of WNK signaling puncta during hypokalemia. These KS-WNK1 dependent foci, termed WNK bodies, influence the spatial localization of the DCT WNK-SPAK/OSR1 pathway^12^ (Fig 1C). Hypokalemic WNK bodies contain WNK4, L-WNK1, SPAK, and OSR1, and their appearance correlates with NCC activation^12,14,15^, suggesting that KS-WNK1’s role in WNK body formation is linked to DCT function.

**Figure 1.**
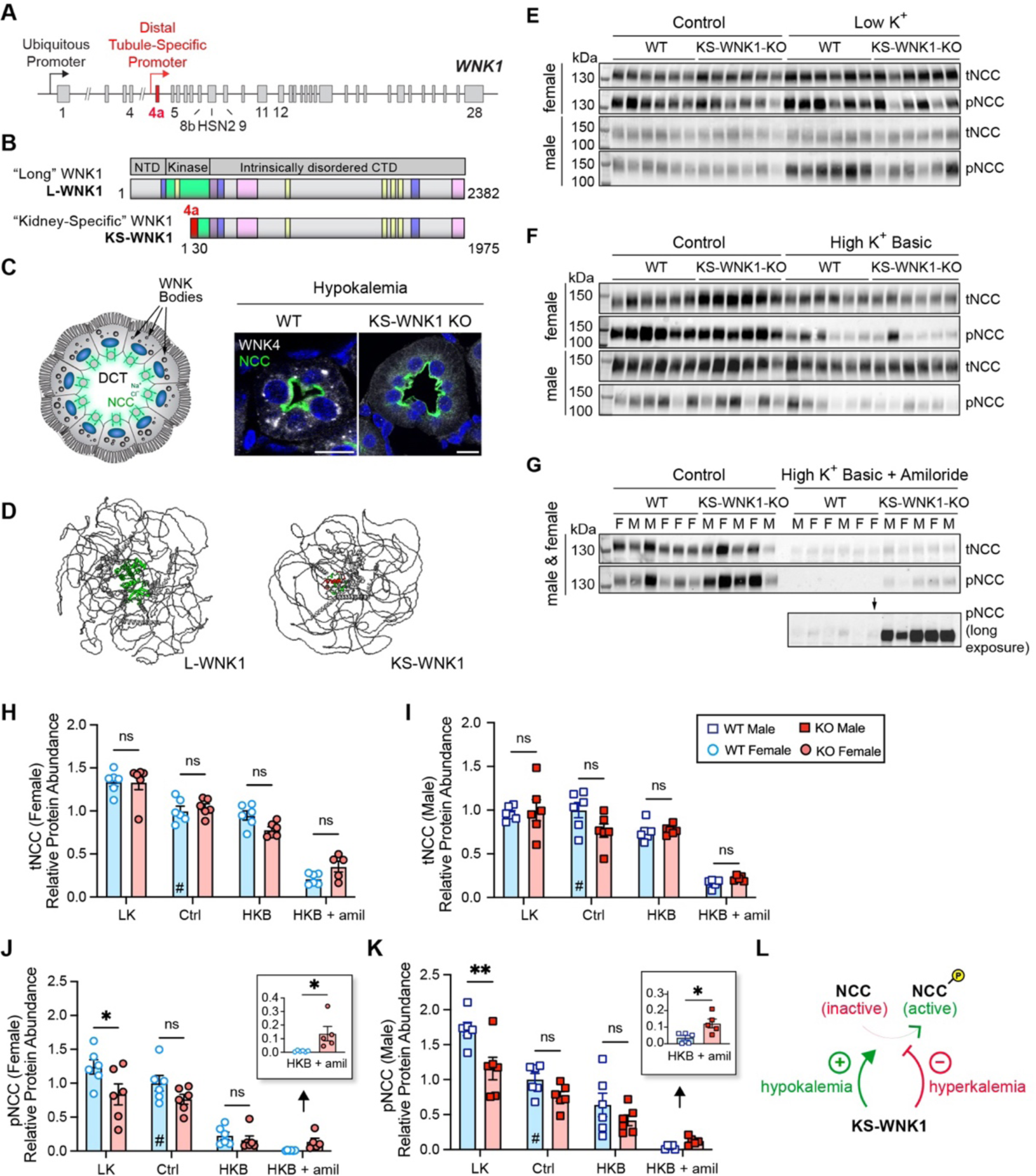
KS-WNK1 differentially alters NCC abundance and phosphorylation during hypo- and hyperkalemia. **A.** Schematic representation of the WNK1 gene. A ubiquitously expressed promoter drives L-WNK1 transcription. A distal tubule-specific promoter drives the expression of the truncated KS-WNK1 isoform. **B.** Top, Domain architecture of L-WNK1 and KS-WNK1 isoforms. Full-length “long” WNK1 contains an N-terminal domain (NTD), a serine-threonine kinase domain (green), and a >100kDa intrinsically disordered C-terminal domain (CTD) that drives condensate formation^18^. Other functional signatures include SPAK/OSR1 binding motifs (yellow), coiled-coil domains (purple), and prion-like regions (pink). KS-WNK1 lacks the L-WNK1 NTD and most of the kinase domain, replaced by a 30AA sequence encoded by exon 4a (red). Schematics adapted with permission from Boyd-Shiwarski et al^12^. **C.** Illustration (left) and immunofluorescence (IF) staining (right) of WNK bodies, which form in NCC-expressing DCT cells during hypokalemia. KS-WNK1 KO mice fail to form WNK bodies. Bar= 10µm. **D.** Alphafold3 predictions of L-WNK1 and KS-WNK1, highlighting their extensive disorder. **E-G.** Immunoblot (IB) analysis of kidney cortical extracts from female and male WT littermates and KS-WNK1 KO mice subjected to 10d maneuvers that alter K^+^ homeostasis. Immunoblots of total NCC (tNCC) and active phospho Thr53 NCC (pNCC), from female and male mice fed either control diet (Ctrl) versus: (E) low K^+^ diet (LK), (F) high K^+^ basic diet (HKB), or (G) high K^+^ basic + amiloride (HKB + amil) (2mg/kg/day). **H-I.** KS-WNK1 had no significant effect on tNCC abundance, regardless of sex. **J-K.** KS-WNK1 had significant effects on pNCC during LK and HKB+amiloride in both male and female mice. Results are shown as means ± SE; *n* = 5-6 mice per genotype per diet. M = male, F = female. Two-way ANOVA with Sidak’s multiple comparisons test, \**P* σ; 0.05, \*\**P* σ; 0.01. **L.** These data indicate that KS-WNK1 stimulates NCC during hypokalemia and inhibits NCC during hyperkalemia. For H-K, data were normalized to WT mice on control diet, indicated by #.

WNK bodies have a unique appearance by electron and light microscopy, as they are micron-sized spherical non-membrane bound regions of the cytoplasm that fail to colocalize with conventional organelle markers^12,16,17^. Thus, they are biomolecular condensates – membraneless cytosolic foci that assemble via phase transitions^18,19^. Though KS-WNK1 dependent WNK bodies are exclusive to the distal nephron, WNK kinases such as L-WNK1 and WNK3 are more ubiquitously expressed, and their ability to form condensates is a fundamental property that allows them to control ion transport and cell volume in many cell types^18,20^. For example, during acute hypertonic cell shrinkage, L-WNK1, SPAK, and OSR1 undergo crowding-induced phase separation, causing their rapid activation within cytosolic liquid-like droplets. Following activation, SPAK and OSR1 leave these dynamic structures and accumulate at the plasma membrane to phosphorylate NKCC1 (*SLC12A2*) and KCCs (*SLC12A4-7*), ubiquitously expressed NCC-like cotransporters that coordinate cell volume recovery. L-WNK1-dependent phase separation is mediated by its large C-terminal domain (CTD), a >100kDa intrinsically disordered region (IDR) that efficiently condenses in response to crowding^18^ (Fig 1B). Notably, the truncated KS-WNK1 isoform lacks intrinsic kinase activity but contains the entire disordered CTD; in fact, this large phase separation-driving IDR comprises greater than 90% of the entire KS-WNK1 protein (Fig 1B,D). This strongly suggests that KS-WNK1 functions as an intrinsically disordered scaffold that coordinates WNK4-SPAK/OSR1 activity in the DCT via condensed phase signaling^21^.

Though KS-WNK1 mediates distal tubule WNK body assembly, its physiologic role in NCC regulation remains unresolved. Here, we report that KS-WNK1 functions as an amplifier of the WNK signaling pathway that operates across the entire physiologic range of [K^+^], allowing the DCT to optimize NCC-mediated salt reabsorption in response to potassium imbalance. By employing mouse models of KS-WNK1 absence and dysfunction, we demonstrate that this effect is WNK body-dependent, establishing a previously unrecognized role for biomolecular condensates in mammalian potassium and blood pressure homeostasis. We further show that the effect is sex-specific, as females require WNK body-mediated signaling to amplify DCT salt reabsorption during hypokalemia. Finally, we show that in the human kidney, WNK body expression progressively increases at a [K^+^] below 4.0mmol/L, suggesting condensate-mediated activation of tubular salt reabsorption when potassium concentrations are within the low-normal reference range. This suggests a role for WNK bodies in human salt-sensitive hypertension. Together, our results identify WNK bodies as kidney-specific signaling condensates that regulate blood pressure and potassium homeostasis.

## Results

### KS-WNK1 amplifies the inverse relationship between NCC phosphorylation and blood [K^+^] via multiple mechanisms

To begin to understand the role of KS-WNK1-dependent WNK bodies in NCC regulation, we studied the effect of KS-WNK1 deletion on NCC expression and phosphorylation status in mice across a broad range of blood potassium concentrations. KS-WNK1-KO mice and wild-type littermates (WT) were administered diets with low K^+^ (LK), control, or alkaline high K^+^ (“high K^+^ basic”; HKB) content for 10 days (Table S1). Because K^+^ loaded mice with a normal glomerular filtration rate efficiently excrete a potassium load^1,22^, we also studied a cohort of HKB-fed mice supplemented with the potassium-sparing diuretic amiloride (2mg/kg/day) to induce frank hyperkalemia (Table S2). We then performed immunoblots for total NCC (tNCC) and phospho-Thr53 NCC (pNCC), a signature of NCC activation (Fig 1E-G, S2A-D).

We initially compared NCC densitometry in WT and KS-WNK1 KO mice stratified by sex and dietary K^+^ maneuver. While this analysis did not reveal a significant effect of KS-WNK1 deletion on tNCC protein abundance (Fig 1H, 1I), KS-WNK1-KO mice exhibited lower pNCC abundance during K^+^ restriction and higher pNCC abundance during K^+^ loading with amiloride (Fig 1E, 1G, 1J, 1K). The effects of HKB with amiloride on pNCC were confirmed with antibodies that recognize alternative NCC phosphoactivation sites at threonine-58 and serine-71 (Fig S2B)^23^. KS-WNK1 KO mice also exhibited higher pNCC abundance than WT controls following 10d of potassium loading on a high KCl (5%) diet (Fig S3A-C), which was sufficient to induce frank hyperkalemia in the absence of a K^+^ sparing diuretic (Fig S3D, S3E). Thus, the effects of KS-WNK1 on pNCC during hyperkalemia were consistent across different physiologic maneuvers. Collectively, these data indicate that KS-WNK1 exerts divergent effects on pNCC depending on potassium status. During hypokalemia, KS-WNK1 appears to be an *activator* of NCC, but during hyperkalemia, it appears to be an NCC *inhibitor* (Fig 1L). These findings comport with a recent report^24^ and suggest that KS-WNK1 exerts complex effects on NCC activity that span the full physiologic spectrum of blood [K^+^].

To further explore these findings, we performed a regression analysis, plotting individual tNCC and pNCC densitometry values for mice subjected to the low K^+^, control, HKB, and HKB + amiloride maneuvers, as a function of blood potassium concentration measured at the time of sacrifice (Fig 2A-C). For WT mice, these graphs revealed a steep increase in pNCC abundance below a potassium concentration of 4mmol/L, indicating a nonlinear relationship between NCC phosphorylation and blood [K^+^]. This amplification effect was evident in the pNCC and pNCC/tNCC graphs, and was clearly blunted in KS-WNK1 KO mice (Fig 2B,C). We first attempted to fit these data to single exponential curves, but this model was inadequate, as it overestimated nearly all of the measured NCC densitometry values at a blood [K^+^] > 6mmol/L for WT mice (Fig 2A-C; compare blue WT curves to filled blue circles). Thus, to stabilize variance in NCC signal across measured blood [K^+^] and better visualize goodness-of-fit, we log-transformed the densitometry data (Fig 2D-F). As predicted by the suboptimal single exponential fits to the untransformed data, simple linear regression of these log-transformations resulted in non-random residuals for WT mice, strongly suggesting that additional components are required to adequately model the observed NCC densitometry dependence on K^+^ (Fig S4A, top graphs).

**Figure 2.**
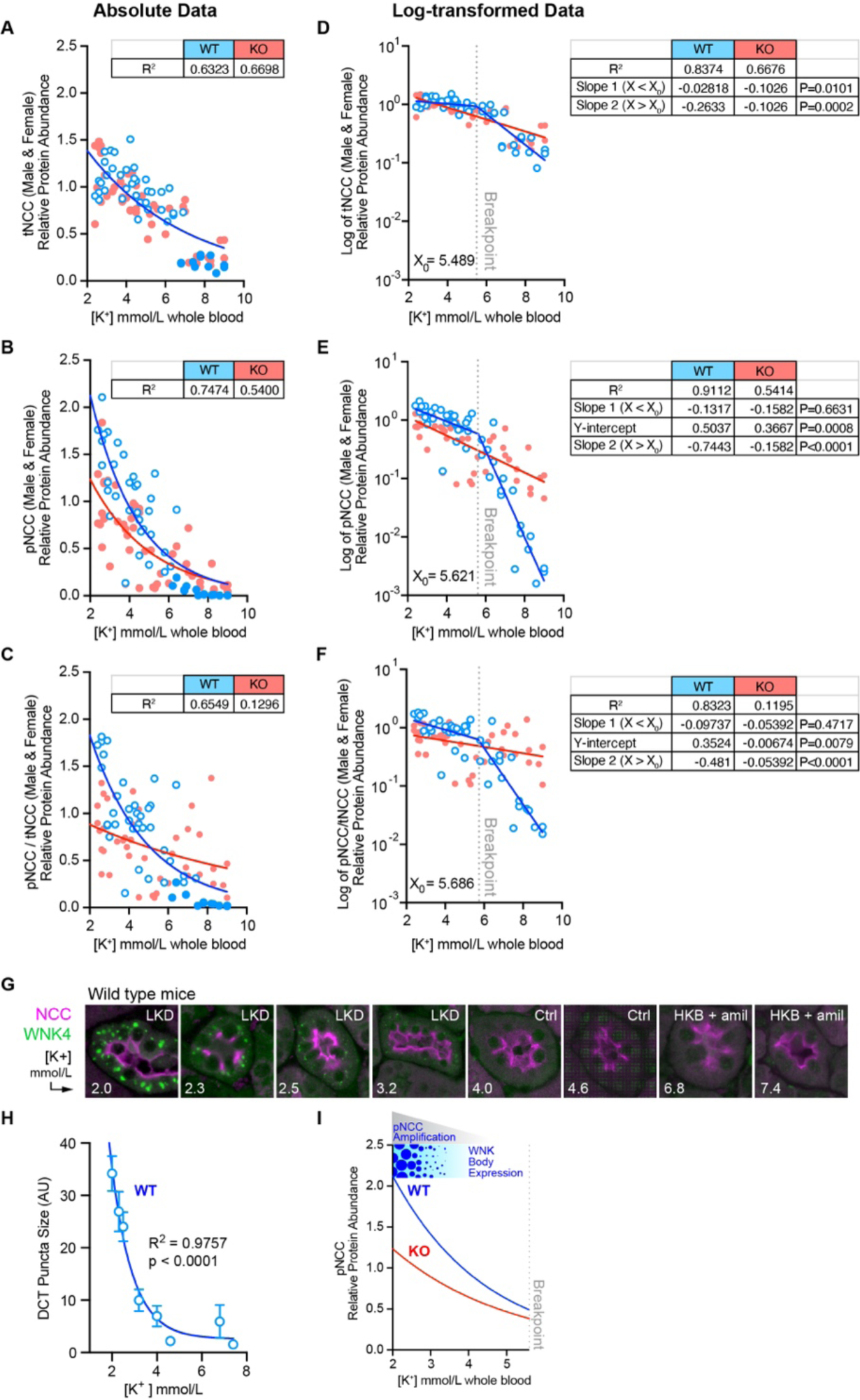
KS-WNK1 amplifies the inverse relationship between NCC phosphorylation and blood [K^+^]. Total and phosphorylated NCC protein abundance in female and male KS-WNK1 KO (red) versus WT (blue) mice, plotted as a function of blood [K^+^]. **A-C**. tNCC, pNCC, and pNCC /tNCC ratio, fit to single exponential curves. R^2^ measures are presented in table format alongside the graphs. For all graphs, the single exponential function adequately fit the WT data at [K^+^] < 4.0, but overestimated data points at [K^+^] >6.0 (filled blue circles). **D-F**. Normalized tNCC, pNCC, and pNCC/tNCC densitometry in A-C were log transformed and analyzed by linear regression. In all cases, WT data were best fit by a segmented linear regression regime, with X0 breakpoints (dotted line) around 5.4-5.6 mmol/L. Slopes of the two linear components are presented in table format alongside the corresponding graphs. For KO mice, Slopes 1 (X<X0) and 2 (X>X0) did not differ as the log-transformed data were best fit by simple linear regression. P-values represent slope comparisons between WT and KO; in the event where Slope 1 comparisons did not reach significance, Y-intercept comparisons with P-values are shown. **G.** IF of DCT WNK bodies in WT male mice treated with various potassium maneuvers for 10d to induce a broad range of blood K+ concentrations, as indicated. DCTs were identified by NCC co-staining. WNK4-positive puncta progressively increased in size as [K^+^] fell and were not visible above a [K+] of 4.0. **H.** Quantification of WNK body size as a function of blood [K^+^], fit to a single exponential curve; R^2^ = 0.9757, p < 0.0001 vs horizontal line (i.e. no dependence on K^+^). **I.** As [K^+^] falls below 4.0mmol/L, WT mice amplify NCC phosphorylation more effectively than KS-WNK1 KO mice, correlating with WNK body expression.

Subsequent curve-fitting trials revealed that the WT log-transformed data were best explained by a *segmental linear regression* model, which added 2 parameters: a second slope and a breakpoint dividing the segments (X0; Fig 2D-F). For WT mice, the addition of a second linear component yielded symmetric and randomly distributed residuals, significantly improving the fit [p ≤ 0.0001 vs straight line by F-test for all three WT curves (tNCC, pNCC, and pNCC/tNCC), Fig S4A bottom]. Remarkably, the tNCC, pNCC, and pNCC/tNCC breakpoints consistently settled at a blood [K^+^] around 5.6 mmol/L; i.e., near the upper limit of the currently defined normal reference range for serum^25^. In contrast to WT mice, the tNCC, pNCC, and pNCC/tNCC data in KS-WNK1 KO mice were best-fit by simple linear regression (Fig 2D-F), as goodness-of-fit was not improved by segmental linear regression. Furthermore, simple linear regression yielded symmetric and randomly distributed residuals for the KO data (Fig S4B). These results indicate that while both WT and KS-WNK1 KO mice exhibit an inverse relationship between NCC and blood [K^+^], the WT mice require an additional component to account for the relationship between NCC and blood [K^+^] during hyperkalemia. These findings were concordant in males and females when the pNCC data were disaggregated by sex (Fig S4C-F).

This regression analysis uncovered key differences in the relationship between potassium and pNCC in WT and KS-WNK1 KO mice. At a [K^+^] less than 5.6 mmol/L (i.e., X<X0), the log-transformed WT and KO slopes of pNCC and pNCC/tNCC were not different, but the Y-intercepts for pNCC and pNCC/tNCC were significantly lower in KS-WNK1 KO mice (p= 0.0008 and 0.0079 for pNCC and pNCC/tNCC respectively; Fig 2E, 2F). Thus, when blood potassium is <5.6 mmol/L, NCC phosphorylation status increases exponentially in both WT and KS-WNK1 KO mice as plasma K^+^ gets progressively lower, an amplification effect that is significantly blunted in mice lacking KS-WNK1. At a blood potassium greater than 5.6mmol/L (X>X0), the inverse relationship between [K^+^] and pNCC increased dramatically in WT mice, resulting in a marked negative deflection in slope (Fig 2E, 2F). This suggests that WT mice recruit an auxiliary process that further suppresses NCC phosphorylation when blood [K^+^] >5.6 mmol/L. Because this deflection in slope is absent in KO mice, the auxiliary dephosphorylation mechanism appears to be KS-WNK1-dependent. Collectively, these data suggest that KS-WNK1 steepens the inverse relationship between blood [K^+^] and NCC activation through discrete mechanisms that differentially affect NCC phosphorylation and dephosphorylation. In other words, KS-WNK1 *expands the dynamic range* of NCC phosphorylation status in response to changes in blood [K^+^], converting small changes in plasma potassium into large effects on pNCC abundance.

### Relationship between blood [K^+^] and WNK body expression

We hypothesized that the potassium-dependent changes in NCC phosphorylation status correspond with altered WNK body expression. To test this, we performed DCT immunostaining for WNK bodies in kidneys harvested from WT male mice with a broad range of blood potassium concentrations. As blood potassium levels progressively decreased below 4 mmol/L, WNK body size increased exponentially (Fig 2G, 2H). By contrast, WNK bodies were not visible in WT males with a plasma potassium greater than 4.0 mmol/L. Thus, similar to the effects of hypokalemia on pNCC abundance, WNK body expression correlates inversely with changes in blood [K^+^] (Fig 2I). This suggests that the nucleation and growth of these condensates is linked to the amplification of NCC activity during potassium deficiency.

### KS-WNK1 increases WNK-SPAK/OSR1 pathway abundance during K^+^ deficiency but does not influence its abundance during K^+^ excess

The DCT-expressed WNK4-SPAK/OSR1 pathway constitutes the canonical NCC activation signal^8^. Given the role of WNK bodies in controlling its localization, we interrogated this signaling cascade in KS-WNK1 KO mice. Consistent with prior reports^2,26^, K^+^ restriction in WT mice upregulated the expression of WNK4, total (t)SPAK, and phospho-(p)SPAK/pOSR1 relative to control diets (Fig 3A, 3D). By comparison, K^+^ restricted KS-WNK1-KO mice exhibited weaker WNK4-SPAK/OSR1 pathway upregulation (Fig 3A, 3D). The low K^+^ mediated increase in pSPAK/pOSR1 likely reflected changes in tSPAK as there was no increase in the pSPAK/tSPAK ratio (Fig 3E). In contrast to the low K^+^ maneuver, WT and KO mice exhibited no differences in WNK4, tSPAK, or pSPAK/pOSR1 expression in the context of HKB or HKB + amiloride treatments that increase blood [K^+^] (Fig 3B, 3C, 3F). Thus, the ability of KS-WNK1 to amplify NCC phosphorylation during K^+^ deficiency correlates with increased WNK4-SPAK/OSR1 signaling via WNK bodies, but its ability to augment NCC dephosphorylation during K^+^ excess does not (Fig 3G).

**Figure 3.**
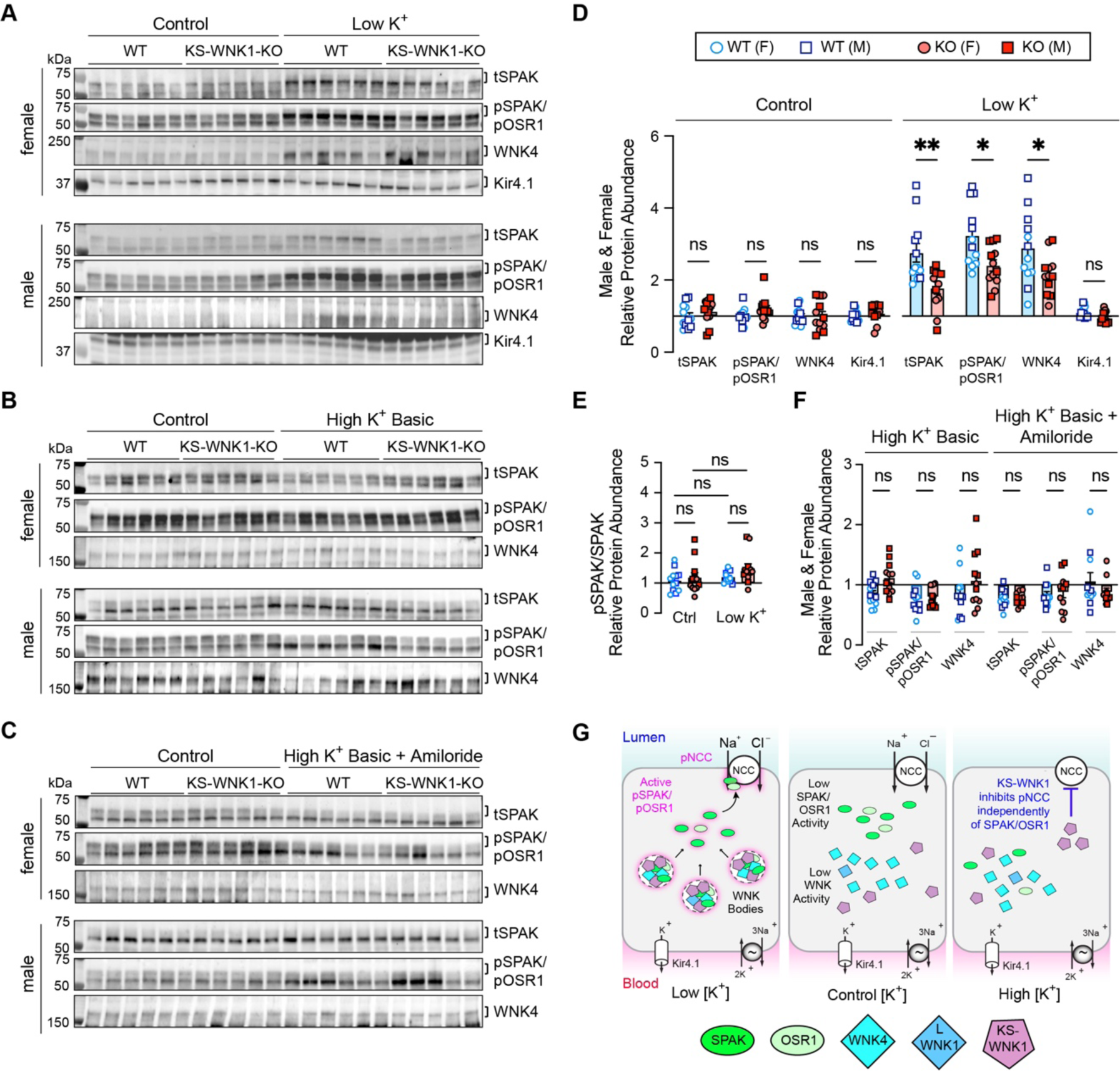
Dysregulated WNK4-SPAK/OSR1 pathway activity in KS-WNK1 KO mice during K+ restriction, but not during K+ loading. IB analysis of kidney cortical extracts from female and male WT littermates and KS-WNK1 KO mice subjected to various K+ maneuvers for 10d. **A-C**. IB of the WNK-SPAK/OSR1 pathway from female and male mice treated with either control diet versus (A) low K^+^ diet, (B) HKB diet, or (C) HKB + amiloride. **D.** WT mice fed a low K^+^ diet had significant increases in tSPAK, pSPAK/pOSR1, and WNK4 compared to WT mice on control diet. KS-WNK1 KO mice had a blunted response to the low K^+^ diet compared to WT mice. **E.** phosphorylated-to-total SPAK ratio in WT and KO mice subjected to control vs low K^+^ diet. **F.** No differences in WNK4-SPAK/OSR1 pathway abundance or phosphorylation in WT and KS-WNK1 KO mice subjected to HKB or HKB + amiloride treatment. **G.** WNK-SPAK/OSR1 pathway localization and activation during low, control, and high blood [K^+^] experimental maneuvers. During low [K^+^], KS-WNK1-dependent WNK bodies condense the WNK-SPAK/OSR1 pathway; this correlates with SPAK/OSR1 and NCC phosphoactivation. During high K^+^, KS-WNK1 inhibits pNCC activation independently of the SPAK/OSR1 pathway. Results are shown as means ± SE; *n* = 6 mice per genotype, sex, and K^+^ maneuver. Two-way ANOVA with Sidak’s multiple comparisons test, \**P* ≤ 0.05, \*\**P* ≤ 0.01.

### KS-WNK1 and WNK body localization

Because the effect of KS-WNK1 on WNK-SPAK/OSR1 signaling likely predominates when hypokalemia-induced WNK bodies are present^12^, we evaluated the effects of KS-WNK1 deletion on WNK-SPAK/OSR1 localization during dietary K^+^ restriction. Consistent with our prior report^12^, K^+^ restriction was associated with the formation of DCT-specific WNK bodies that stained positive for WNK1, WNK4, and pSPAK/pOSR1, suggesting that WNK4 activates its downstream targets within condensates (Fig 4A). These structures were largely absent in potassium-deficient KS-WNK1-KO mice regardless of sex (Fig 4A & D). Even so, pSPAK/pOSR1 apical staining was present despite KS-WNK1 deletion (Fig 4B), indicating that KS-WNK1 is not necessary for active SPAK to engage with NCC during hypokalemia. However, given the observation that KS-WNK1 KO mice exhibit blunted SPAK/OSR1 and NCC phosphorylation during K^+^ deficiency (Fig 1E, 1J, 1K, 2B, 2E, 3A, 3D), the data indicate that KS-WNK1 potentiates SPAK/OSR1 activation in response to decreased blood [K^+^], likely through WNK body-mediated signaling.

As reported previously^12^, KS-WNK1-KO DCT cells rarely exhibited WNK-SPAK/OSR1 positive puncta which were mislocalized to the basal pole during K^+^ restriction (Fig 4B, arrowheads). Morphometric analysis (Fig 4C) revealed that while both sexes exhibited the same number of WNK bodies per cell during K^+^ restriction (Fig 4D), females exhibited larger condensates that were positioned closer to the tubular lumen (Fig 4E & F). These findings suggest sex-specific differences in WNK-SPAK/OSR1 pathway functionality.

**Figure 4.**
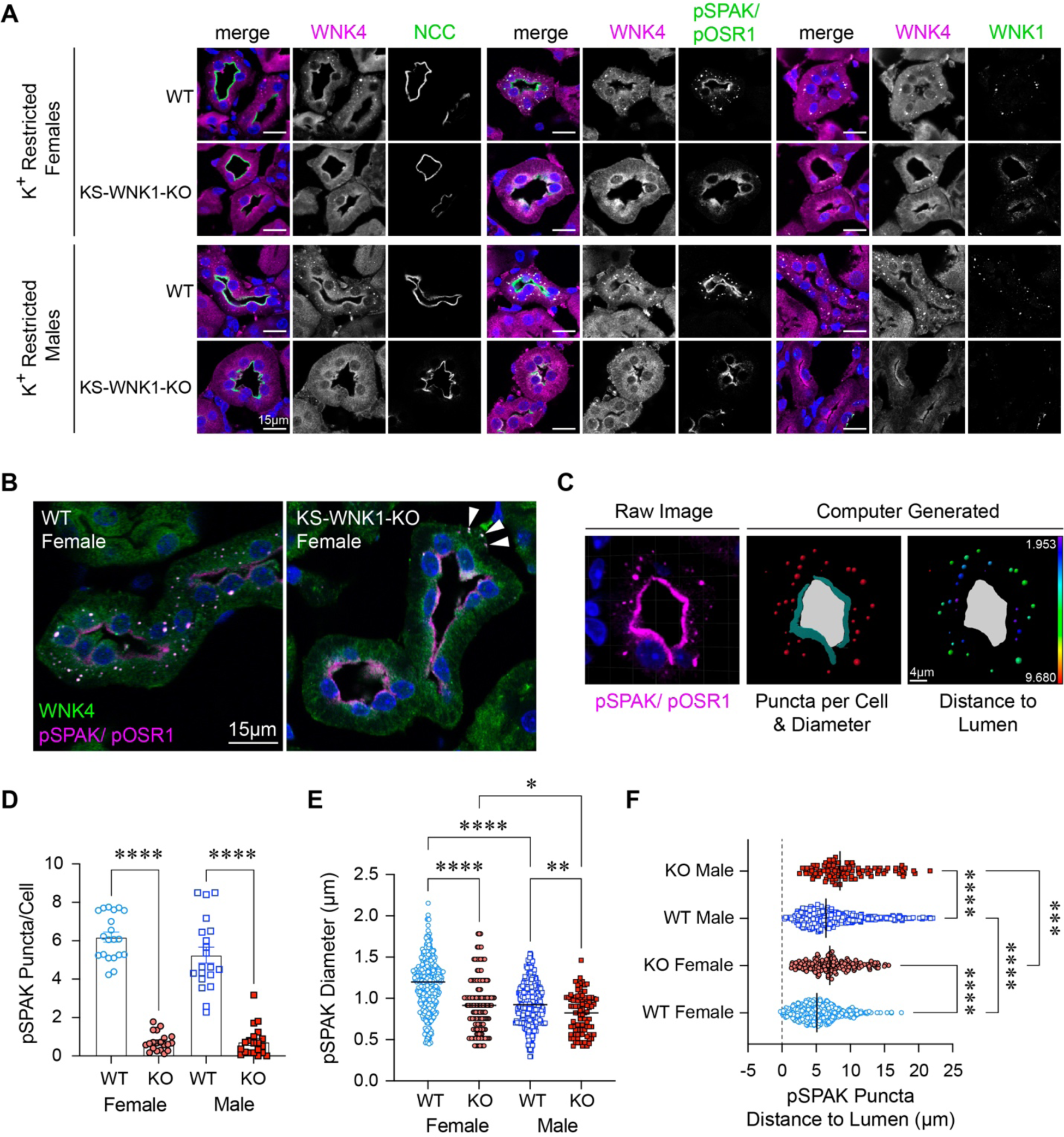
K^+^ restricted WT and KS-WNK1 KO mice exhibit sex differences in WNK body expression. **A.** IF of kidney sections from WT or KS-WNK1 KO mice treated with low K^+^ diet for 10 days. DCTs were identified by NCC co-staining and morphology. WNK4, pSPAK/pOSR1, and WNK1 antibodies colocalized within puncta in WT mice, whereas puncta were nearly absent in KS-WNK1 KO mice. All bars = 15µm. **B.** WNK body formation in female WT and KS-WNK1 KO mice. Cytosolic puncta are largely absent in KS-WNK1-KO mice, though apical pSPAK/pOSR1 staining is still present. Uncommonly, mislocalized basolateral puncta containing pSPAK/pOSR1 and WNK4 were observed (arrowheads). Bar = 15µm. **C.** Imaris was used to quantify WNK body number and size (*middle*) and distance to lumen (*right*), from raw confocal IF images of pSPAK/pOSR1 intracellular puncta in DCTs (*left*). Bar = 4µm. **D-F.** Quantification of pSPAK/pOSR1 (D) Puncta per cell (20 tubules per condition), (E) Puncta diameter (5 tubules per condition), and (F) Distance to apical lumen in female and male mice (5 tubules per condition). Two-way ANOVA with Sidak’s multiple comparison, * P ≤ 0.05, ** P ≤ 0.01, *** P ≤ 0.001, **** P ≤ 0.0001.

### Sex-specific effects of KS-WNK1 on blood and urine electrolytes

We also observed diverging trends between WT and KS-WNK1 KO mice when blood and urine data were disaggregated by sex. Sex-dependent differences were more evident at the extremes of low and high K^+^ and affected blood K^+^, Cl^−^, HCO3^−^, Ca^2+^, and urine osmolarity and pH. When placed on a K^+^ deficient diet, where NCC phosphorylation is high and KS-WNK1 dependent (Fig. 1), KS-WNK1 KO females exhibited more pronounced hypernatremia, hypokalemia, and reduced urine osmolality compared to WT females (Fig 5A-C, Table S2 & S3). We did not observe differences in urine [K^+^] as these measurements were at the low limit of detection on a K^+^ deficient diet (Table S3). In contrast to females, KS-WNK1 KO males did not exhibit lower blood [K^+^] versus WT males during potassium restriction (Fig 5C), despite strong effects of KS-WNK1 deletion on pNCC abundance (Fig 1). Male KS-WNK1 KO mice also did not exhibit differences in blood [Na^+^] or urine osmolality. Relative to WT littermates, however, male KS-WNK1 KO mice were more hypercalcemic during K^+^ restriction (Fig 5D, Table S2). Thus, K^+^-restricted KS-WNK1 KO mice exhibit features commonly seen in states of low NCC activity, such as Gitelman syndrome or thiazide diuretic administration^8^.

**Figure 5.**
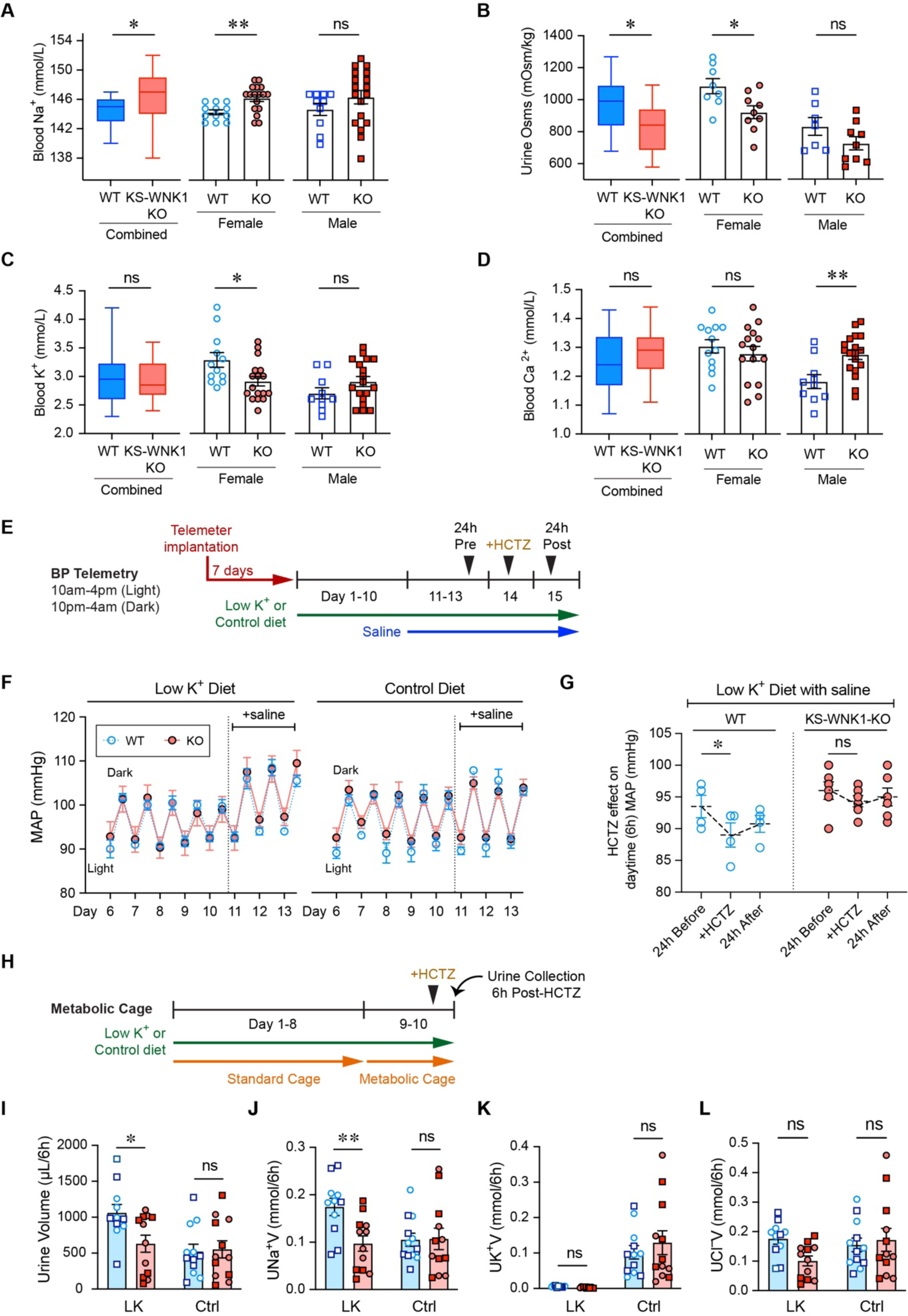
Effect KS-WNK1 on blood and urine composition, and blood pressure thiazide-sensitivity in K^+^ restricted female and male mice. Male and female WT and KS-WNK1 KO mice were fed low K^+^ diets for 10 days and whole blood electrolytes and urine were obtained on day 10. Results were analyzed as both combined and sex-disaggregated data. **A.** KS-WNK1 KO mice had significantly increased blood [Na^+^] when males and females were analyzed in combination, and when females were analyzed separately. **B,** Urine osmolality in KS-WNK1 KO mice was significantly decreased in the combined male and female dataset, and in the female pool. Mean Uosm in KS-WNK1 KO males was lower than male WT without reaching significance. **C.** KS-WNK1 deletion had no significant effect on whole blood [K^+^] in the combined pool, or in the male pool. However, female KS-WNK1-KO mice had a significant decrease in whole blood [K^+^] compared to sex-matched controls. **D.** KS-WNK1 deletion had no significant effect on whole blood [Ca^2+^] in the combined male and female pool or in the female pool. However, male KS-WNK1-KO mice had a significant increase in whole blood [Ca^2+^]. Sample size: n= 10-18 mice. Unpaired t-test between WT and KO was used to determine significance, \**P* ≤ 0.05, \*\**P* ≤ 0.01. **E.** Schematic of the BP telemetry experiment. Female WT (*n* = 4) and KS-WNK1 KO mice (*n* = 6) were subjected to either low K+ or control diets for 10 days, followed by supplementation with 1% normal saline (NS) in drinking water for 3 days. **F.** KS-WNK1 expression had no significant effect on mean arterial pressure (MAP) in K^+^ restricted or control diet mice. Saline supplementation increased MAP in K^+^ restricted mice. Genotype had no significant effect based on two-way ANOVA with post-hoc Sidak’s multiple comparison test. Each data point represents either daytime or nighttime MAP collected over 6h. **G.** Thiazide diuretic challenge. Mice were fed low K+ diet for 10 days and then given 1% saline in their drinking water for 3 days. Daytime MAP was measured for a 6h window starting 24 h before HCTZ injection, 1h after HCTZ injection (+HCTZ, 25mg/kg IP) and 24 h after HCTZ injection. WT mice had a significant decrease in MAP with HCTZ treatment, whereas KS-WNK1 KO mice had a blunted response to HCTZ. *P ≤ 0.05, two-way ANOVA with post-hoc Sidak’s multiple comparisons test. **H.** Schematic of the metabolic cage experiment. Diuretic challenge was performed in both female and male WT and KS-WNK1 KO mice. After 10 days of control or low K^+^ diet, mice were injected with HCTZ (25mg/kg IP), and urine was collected over 6h. **I-L.** (I) urine volume and (J) urine Na*V (UNaV) in were greater in WT mice compared to KO mice. (K) On the low K^+^ diet, urine K^+^ was too low to detect a significant difference. (L) There was a trend for HCTZ to blunt Cl-excretion in KS-WNK1 KO mice on low K+ diet, without reaching significance. Results are shown as means ± SE; *n* = 5-6 mice per genotype, sex, and diet. Two-way ANOVA with Sidak’s multiple comparisons test; \**P* ≤ 0.05, \*\**P* ≤ 0.01.

Next, we investigated whether the relative hypokalemia in K^+^ restricted KS-WNK1-KO females was due to changes in aldosterone, ENaC, or ROMK – known factors that can cause hypokalemia. If these factors were driving relative hypokalemia in KO mice, these mice would be expected to have higher aldosterone levels and increased ENaC and ROMK abundance; however, that is not what was observed. During dietary K^+^ restriction, aldosterone (aldo) levels tended to be lower in KS-WNK1 KO mice (Table S2), consistent with prior observations^27^. We did observe lower expression of the uncleaved form of γENaC (Fig S5A-C) and could not detect expression of the subunit’s cleaved/active form in either WT or KO mice during dietary K^+^ restriction (Fig S5A-C). We also found that ROMK protein abundance was not elevated in KS-WNK1-KO female mice relative to WT sex-matched littermates (Fig S5D, S5E). Collectively, these findings suggest that the hypokalemia in female KS-WNK1 KO mice is not due to increased aldosterone, ENaC, or ROMK, but instead reduced NCC activity.

### Role of KS-WNK1 in blood pressure regulation, salt sensitivity, and thiazide responsiveness

Given the importance of NCC in blood pressure regulation^3^, we performed telemetric blood pressure measurements in KS-WNK1 KO mice (Fig 5E). These studies focused on females since they exhibited larger electrolyte differences during K^+^ restriction (Fig 5A-C). Despite differences in pNCC expression (Fig 1), ten days of potassium deprivation yielded no differences in mean arterial pressure (MAP) between WT and KS-WNK1 KO females (Fig 5F). Moreover, though K^+^ restricted WT and KO mice both developed a salt-sensitive increase in blood pressure^22^, we observed no differences between knockout mice and WT littermates. Female mice administered a control diet for 10d did not exhibit changes in blood pressure, either at baseline or after saline-loading, regardless of KS-WNK1 genotype (Fig 5F). To test for differences in NCC activity, we challenged potassium restricted, saline-loaded female WT and KS-WNK1 KO mice with an intraperitoneal injection of hydrochlorothiazide (HCTZ; 25mg/kg). WT mice responded to HCTZ injection with a significant 4.5 mmHg average decrease in MAP. In contrast, KS-WNK1 KO mice were relatively insensitive to HCTZ—(Fig 5G). Thus, similar to humans with Gitelman syndrome^28^, K^+^ restricted KS-WNK1 KO females exhibit low NCC activity but are able to maintain their blood pressure via compensatory effects.

To further assess the effects of KS-WNK1 on NCC activation, we administered HCTZ and measured urinary volume, [Na^+^], [K^+^], and [Cl^-^] in male and female mice maintained on either a low K^+^ or control diet for 10 days (Fig 5H). K^+^ restricted KS-WNK1 KO mice exhibited a blunted response to HCTZ compared to WT littermates, with decreased urinary volume and UNa^+^V (Fig 5I, 5J). There was no change in urinary K^+^, likely because it was near the lower limit of detection due to dietary K^+^ restriction (Fig 5K). There was also a trend for decreased urinary [Cl^-^] under K^+^ restricted conditions that did not reach significance (Fig 5L). These diuretic and natriuretic responses reflect lower NCC activity in KS-WNK1 KO mice compared to WT controls, consistent with the reduced NCC phosphorylation in KS-WNK1 KO mice after dietary K^+^ restriction (Fig 1 & 2).

### WNK bodies are required for NCC activation during hypokalemia

Our analyses in knockout mice demonstrate that during potassium deficiency, KS-WNK1 drives WNK body formation (Fig. 4), amplifies NCC phosphoactivation (Fig 1), and increases WNK4-SPAK/OSR1 pathway expression (Fig 3). However, these studies do not establish whether the blunted NCC activation in K^+^ restricted KS-WNK1 KO mice is specifically due to impaired WNK body condensation or other KS-WNK1 dependent factors. To address this, we generated a mouse expressing a full-length mutant version of KS-WNK1 that cannot form functional WNK bodies. As noted in Fig 1, the KS-WNK1 N-terminus is capped by 30 unique amino acids encoded by exon 4a. Previously, we showed that the exon 4a coding sequence contains a conserved cysteine-rich hydrophobic (CRH) motif that is necessary for WNK body formation in vitro^12^. To disrupt WNK body formation in mice, we replaced five essential hydrophobic residues within the CRH motif with five neutral glutamines (VFVIV->QQQQQ) (Fig 6A, S6A-D). AlphaFold predicts this “5Q” mutation disrupts KS-WNK1 N-terminal structure, unraveling an amphipathic helix encoded by exon 4a (Fig 6B).

**Figure 6.**
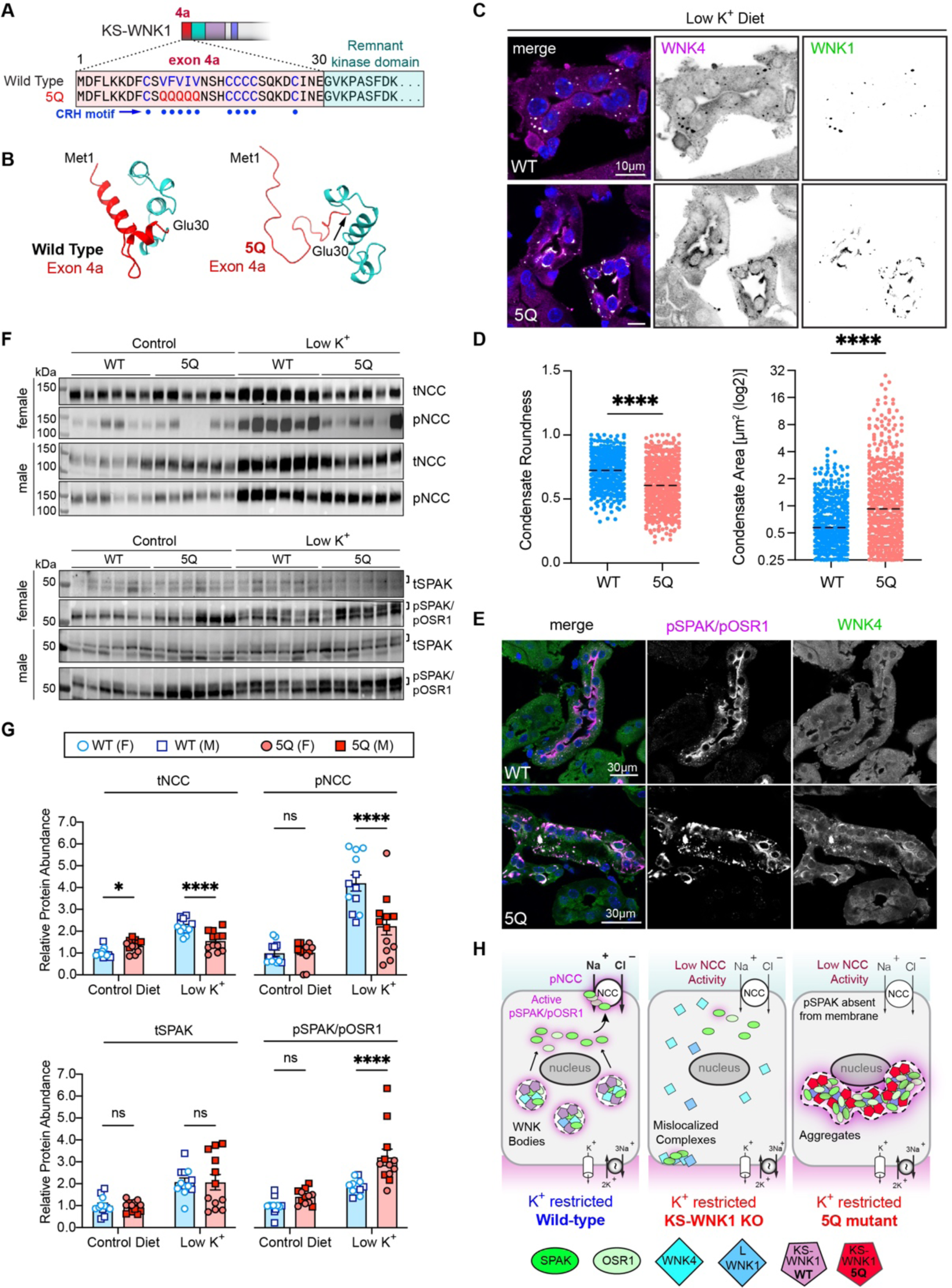
WNK bodies are necessary for KS-WNK1 to amplify NCC phosphorylation during hypokalemia. **A.** Exon 4a of KS-WNK1 encodes 30 amino acids, including a cysteine-rich hydrophobic (CRH) motif. The motif’s patch of 5 consecutive bulky hydrophobic residues was mutated to 5 glutamines to generate “5Q” mice with aberrant WNK body formation. **B.** AlphaFold predicted structures of the WT and 5Q exon 4a peptide (red) and the adjacent remnant kinase domain (cyan). The 5Q mutation disrupts a predicted helical structure encoded by exon 4a. **C.** IF of kidneys from female 5Q mice maintained on low K^+^ diet for 10 days. Typical WNK bodies are absent and replaced by irregularly shaped foci that often form paranuclear crescents and contain WNK4 and pSPAK/pOSR1. **D.** Quantification of WNK condensate morphology in WT and 5Q mice. The 5Q foci were less round and larger than wild-type WNK bodies. N=425 objects from 9 images for WT, 530 from 12 images for 5Q, \*\*\*\**P* < 0.0001. **E.** WNK4 and pSPAK/pOSR1 costaining in WT and 5Q mice. In WT, pSPAK/pOSR1 signal colocalized with WNK4 in puncta and was also located at the DCT apical membrane. In contrast 5Q mice exhibited strong pSPAK/OSR1 and WNK4 co-condensation, but no apical pSPAK/pOSR1. **F.** WT and 5Q mice were fed control or low K^+^ diet for 10d and kidney cortex homogenates were probed for tSPAK, pSPAK/pOSR1, tNCC, and pNCC. **G.** graphical representation of immunoblots in (F). K^+^ restricted 5Q mice had significantly increased pSPAK/pOSR1 and reduced tNCC and pNCC expression, indicating that signaling to NCC was uncoupled. *n* = 6 mice per genotype per diet. Two-way ANOVA with Sidak’s multiple comparisons test, \**P* ≤ 0.05, \*\**P* ≤ 0.01. **H.** Model of WNK-SPAK/OSR1-NCC signaling in WT, KS-WNK1 KO, and KS-WNK1 5Q mice. KS-WNK1 normally facilitates WNK body condensate formation and NCC activation via the WNK-SPAK/OSR1 pathway. In K^+^ restricted KS-WNK1 KO mice, WNK bodies are largely absent and remaining complexes are mislocalized, resulting in low SPAK/OSR1 and NCC activity. In the K^+^ restricted 5Q mouse, WNK-pSPAK/pOSR1 becomes trapped in aggregates that surround the nucleus preventing pSPAK/pOSR1 expression at the DCT apical membrane, causing a reduction in NCC activity.

Mice homozygous for the KS-WNK1 5Q mutation were maintained on control or low K^+^ diet for 10d and assessed for changes in SPAK/OSR1 and NCC activation by immunostaining, western blot, and electrolyte measurements. Potassium-restricted 5Q mice were unable to form spherical WNK bodies, and instead formed larger amorphous aggregates with reduced roundness that accumulated around nuclei (Fig 6C, 6D). These structures were enriched in pSPAK/pOSR1, which was absent from the apical membrane suggesting aberrant function (Fig 6E). Consistent with intracellular sequestration and disruption of signaling, during K^+^ restriction pSPAK/pOSR1 abundance was increased in kidney immunoblots by 210% in males (p = 0.0019) and 136% in females (p = 0.0032), but pNCC abundance did not increase accordingly. Instead, pNCC decreased by 25% in males (p =0.018) and 63% in females (p = 0.0008) (Fig 6F, 6G, S7C-F).

Similar to KS-WNK1 KO mice, potassium-restricted female 5Q mice had a significantly decreased blood [K^+^] and [Cl^-^], and increased [HCO3^-^], suggestive of a sex-specific Gitelman-like phenotype (Table S4). On control diet, 5Q males exhibited hyperchloremic metabolic acidosis and females trended toward this, with elevated [Cl-] and lower [HCO_3_^-^]. Collectively these findings indicate that the KS-WNK1 5Q mutation alters WNK body formation and function during K^+^ restriction, resulting in mislocalization of the WNK-SPAK/OSR1 pathway and low NCC activity. While the 5Q mice can activate DCT-expressed SPAK and OSR1, the phosphorylated forms of these kinases are trapped within dysfunctional cytoplasmic aggregates and thus are unable to activate NCC at the apical membrane. Therefore, functional WNK bodies are necessary for optimal NCC activation during hypokalemia.

### WNK body abundance correlates with serum [K^+^] in humans

In WT mouse models, WNK bodies form during hypokalemia and disperse during normokalemia^12,14^ (Fig 2G). To date, human WNK bodies have only been reported in the setting of hypokalemic nephropathy, a pathologic condition caused by severe K^+^ deficiency^17^. To explore the physiological relevance of WNK bodies in human health, we asked whether these potassium-dependent condensates are present in humans with [K^+^] in the physiologic range (3.5-4.2 mmol/L). WNK bodies were present in all 6 kidney samples studied from both male and female subjects ages 46-79 years (Fig 7A). Consistent with studies in mice, there was an inverse correlation between decreasing serum [K^+^] and increasing WNK body abundance (Fig 7B, 7C). A progressive increase in WNK bodies per DCT cell was noted, particularly when the measured [K^+^] was <4.0mmol/L (Fig 7B). This suggests that in humans, WNK bodies activate NCC via the WNK-SPAK/OSR1 pathway, even when blood potassium concentrations are in the low-normal reference range of 3.5-4.0 mmol/L.

**Figure 7.**
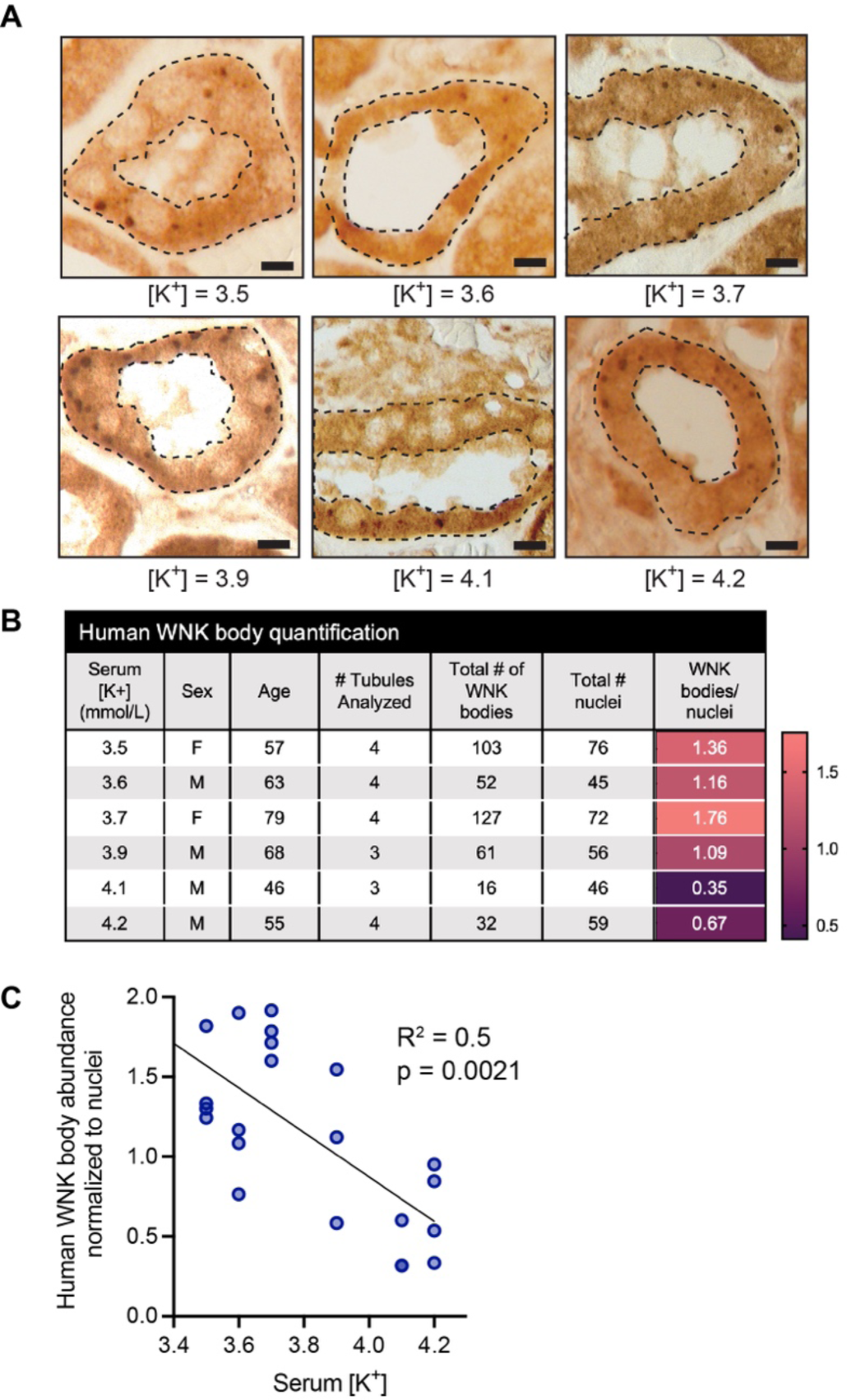
Human WNK body abundance correlates with serum [K^+^]. **A.** Immunohistochemistry of distal convoluted tubules obtained from 6 human kidney wedge biopsies stained for WNK1. DCTs were confirmed by NCC staining in adjacent sections (not shown). **B.** Serum [K^+^], sex, and age of the subjects, along with the values used for quantification. A heat map indicates the correlation between WNK bodies with increasing serum [K^+^]. **C.** There was an inverse relationship between serum [K^+^] and WNK body abundance. To calculate the number of WNK bodies per cell, the number of WNK bodies within a single tubule were counted and then normalized to the number of nuclei within that tubule. Results are shown as means ± SE; *n* = 3-4 tubules analyzed per subject. R^2^ = 0.5, p < 0.0021 vs horizontal line (i.e. no dependence on K^+^).

## Discussion

First reported over a decade ago, WNK bodies were initially described as punctate clusters of the WNK-SPAK/OSR1 pathway that form in the DCT during potassium deficiency^15,29^. Subsequent studies noted that these foci are membraneless, consistent with the notion that they are specialized biomolecular condensates that assemble during potassium stress^12,19^. WNK body formation requires KS-WNK1, but the role of this isoform in DCT salt transport has been elusive: several studies have suggested that KS-WNK1 inhibits NCC-mediated sodium transport^27,30–33^, while others claim that it is an NCC activator^34–36^. These apparent contradictions are reconciled when the effect of KS-WNK1 on pNCC is analyzed as a function of blood potassium concentration. Subjecting over 100 KS-WNK1 KO mice and littermate controls to various potassium maneuvers designed to manipulate NCC phosphorylation across a wide physiologic range, we found that KS-WNK1 KO mice exhibit reduced pNCC during hypokalemia and increased pNCC relative to WT mice during hyperkalemia. Thus, KS-WNK1 functions as both an activator and an inhibitor of NCC, depending on potassium status. Integrating these results as a function of blood [K^+^] in regression analyses, we found that the steep inverse relationship between NCC phosphorylation and potassium is blunted in KS-WNK1 KO mice. This indicates that KS-WNK1’s true function is to expand the dynamic range of NCC phosphorylation across the entire physiologic spectrum of blood potassium concentrations that are experienced during life. In order for the DCT to adjust salt reabsorption in response to potassium imbalance, it must sense small fluctuations in interstitial potassium concentrations and then convert those tiny changes into robust effects on NCC activity. Our findings demonstrate that KS-WNK1 is an essential part of this signal amplification mechanism.

Though KS-WNK1 augments NCC phosphorylation during hypokalemia and dampens NCC phosphorylation during hyperkalemia, the mechanisms of action are distinct. During K^+^ loading, the relationship between KS-WNK1 and pNCC change as the blood potassium concentration rises above 5.6 mmol/L. Above this breakpoint, pNCC abundance decreases dramatically, resulting in a steep negative deflection in slope. Since this feature is not present in KS-WNK1 KO mice, our findings suggest that KS-WNK1 recruits an auxiliary mechanism that promotes NCC dephosphorylation during hyperkalemia, independently of WNK4-SPAK/OSR1 signaling. The process also appears to be WNK body-independent, as these condensates are not visible in hyperkalemic WT mice. Though the underlying mechanism is not clear, our data imply that KS-WNK1 promotes the action of NCC-specific phosphatases, especially given their established role in NCC dephosphorylation during hyperkalemia^4^.

Studies in the KS-WNK1 5Q mutant mouse demonstrate that the mechanism by which KS-WNK1 amplifies WNK-dependent signaling during hypokalemia is condensate-dependent. During hypokalemia, KS-WNK1 recruits the WNK-SPAK/OSR1 pathway within WNK bodies to facilitate efficient activation. Similar to other biomolecular condensates^37,38^, a major function of KS-WNK1-dependent WNK bodies is to organize the WNK pathway within hubs that coordinate condensed phase signaling; in this case, to amplify changes in distal tubule sodium chloride reabsorption during hypokalemia. Though the molecular basis by which WNK bodies augment WNK-SPAK/OSR1 pathway activity remains unresolved, it appears that normal KS-WNK1 exon 4a structure is required to coordinate the condensed NCC amplification signal. Mutagenic disruption of exon 4a results in the formation of dysfunctional aggregates that trap the WNK-SPAK/OSR1 pathway. This strongly suggests that WNK bodies i) possess a defined structure that scaffolds and organizes WNK4-dependent signaling complexes, ii) facilitate efficient SPAK and OSR1 activation, and iii) allow the activated kinases to leave the condensed phase and traffic to the DCT apical membrane so they can phosphorylate NCC. Thus, WNK bodies represent a clear example of a specialized stress-induced biomolecular condensate that physiologically regulates whole body electrolyte homeostasis via signal amplification.

Our results also identify KS-WNK1-dependent WNK bodies as key determinants of sex-based differences in distal nephron function. Our data suggest that female mice require DCT WNK bodies to minimize salt wasting and maintain potassium homeostasis. Recent observations indicate that females prioritize distal tubule salt reabsorption via NCC more than males^39,40^. The reason for this is unclear, but a role for NCC in defending against hypokalemia during pregnancy has been proposed^41^. Regardless, our data suggest that the underlying mechanism is dependent on the ability of females to leverage the distal tubule and WNK body-mediated signaling more efficiently during the threat of potassium deficiency. Given the influence on whole animal physiology, our findings more generally imply that biomolecular condensates may contribute to sexual dimorphism in other regulatory systems.

Lastly, our results implicate WNK bodies as regulators of potassium homeostasis in humans. The human kidney evolved to conserve electrolytes and blood volume during times of undernourishment, and to tolerate the diets consumed by our paleolithic ancestors during times of surfeit. The paleolithic diet, rich in K^+^ and low in Na^+^, differs greatly from the high salt/ low K^+^ diets that are consumed by most people today^42^. The current average US dietary intake of potassium is well below recommended guidelines^43^, and low K^+^ intake has been associated with an increased incidence of hypertension, cardiovascular disease, and all-cause mortality^44^. This suggests that modern “Western” diets may activate tubular stress responses that preserve [K^+^] at the expense of excessive sodium chloride reabsorption and blood pressure salt-sensitivity. In normal human kidney parenchyma, we found an inverse relationship between WNK body condensate abundance and plasma potassium that progressively increased as [K^+^] fell below 4.0 mmol/L. Thus, humans with normal kidney function appear to exhibit hyperactive WNK signaling and distal salt reabsorption at the low end of the normal reference range for [K^+^]. This raises an intriguing question about the current laboratory definition of normokalemia: Is a [K^+^] of 3.5 mmol/L truly “normal” if it predisposes to salt-sensitive hypertension? Indeed, a recent meta-analysis comprising greater than 1 million participants found that the risk of adverse cardiovascular outcomes, end stage kidney disease, and all-cause mortality increases as blood [K^+^] falls below 4.0 mmol/L^45^. From a therapeutic standpoint, our findings support the concept that potassium supplementation can control blood pressure, as has been touted in recent studies and current hypertension guidelines^44,46,47^. Conceivably, titrating K^+^ intake to achieve a plasma potassium concentration sufficient to inhibit WNK body condensation may be an effective antihypertensive strategy in certain clinical scenarios.

In conclusion, our findings provide insight into the role of KS-WNK1 and WNK bodies in potassium-dependent NCC regulation in both mice and humans. They identify KS-WNK1 as a DCT-specific amplifier that optimizes NCC reactivity to changes in systemic potassium concentrations. This observation reconciles conflicting data regarding stimulatory vs inhibitory effects of KS-WNK1 on NCC. Key to the amplification process is KS-WNK1’s ability to organize the WNK signaling pathway within specialized biomolecular condensates that impact potassium metabolism, blood pressure salt sensitivity, and nephron sexual dimorphism. WNK body assembly requires KS-WNK1, which emerged during vertebrate kidney evolution from a crowding-sensitive condensation-prone kinase that first appeared in single celled organisms to control salt transport and fluid volume^18,19^. Thus, it appears that through nephron segment-specific isoform expression, the mammalian kidney repurposed this ancient condensate-dependent volume regulatory system for total body potassium and blood pressure homeostasis.

## Acknowledgments

We thank Tom Kleyman for helpful discussions. This content is solely the responsibility of authors and does not necessarily represent the views of the U.S. Department of Veterans Affairs.

## Author Contributions

C.B.S. and A.R.S. designed the study; C.B.S, R.T.B., J.A.L., M.N.V., A.B, S.A.K., S.E.G., L.J.N., K.Q., A.L.M., and S.D.S. performed experiments; C.B.S., D.J.S., M.N.V., N.H.N, A.B, S.D.S., S.A.K., O.B.K., and A.R.S. analyzed data; C.B.S., D.J.S., and A.R.S. made figures; C.B.S., R.T.B., S.E.G., D.J.S., A.R.R., S.D.S., O.B.K., and A.R.S. drafted the paper; C-L.H. provided KS-WNK1 knockout mice; all authors approved the final version of the manuscript.

## Methods

### Mice

All animal protocols conform to the National Institutes of Health (NIH) Guide for the Care and Use of Laboratory Animals and were approved by the University of Pittsburgh IACUC. KS-WNK1 knockout mice (KS-WNK1-KO) and age-matched wild-type littermates (WT) were generated in a 129/Sv background as previously described ^12,31^. KS-WNK1-KO mice were derived from a mouse line originally reported by Liu *et al*. ^31^ and described in the supplemental materials from Boyd-Shiwarski *et al.* ^12^. Genotyping was performed as reported by these studies previously. CRISPR-Cas9 homology-directed repair was used to knock-in a mutation in exon 4a of the WNK1 gene, replacing five consecutive bulky hydrophobic residues (spanning Val-11 to Val-15) with five neutral glutamines, resulting in the generation of KS-WNK1 “5Q” mutant mice (Fig 6A, S6A-D). The 5Q mutant mice were generated and bred in a 129/Sv background. Experiments were performed on both female (20-30g) and male mice (25-35g), aged 10-25 weeks. All mice were housed in a temperature-controlled room on a 12h light/dark cycle. Mice had free access to deionized water, unless otherwise noted.

### Dietary Maneuvers

To determine the effect of KS-WNK1 on NCC phosphorylation, mice were fed K^+^ diets for 10 days: (1) low K^+^ (LK, TD.88239), (2) control K^+^ (ctrl, TD.88238), (3) high K^+^ basic, (HKB, TD.07278), (4) high KCl, (HKCl, TD.09075) (Teklad, Madison, WI). See Table S1 for Teklad Diet concentrations of the following minerals: Na^+^, K^+^, Cl^-^, Ca^2+^. To induce hyperkalemia, mice fed the high K^+^ basic diet were supplemented with amiloride (2mg/kg/d) in their drinking water for 10 days. After 10 days, mice were anesthetized with isoflurane, and blood was obtained via terminal cardiac puncture and analyzed by iSTAT (Abbot). All mice had access to food and water until time of sacrifice and sacrificed the same time of day (between 1-3pm). Kidneys were harvested and flash frozen for immunoblot and/or paraformaldehyde-treated for microscopy. Urine was immediately collected from mouse bladder for urine pH measurements using AimStrip US-5 (Germaine Inc, San Antonio, TX). Blood plasma was isolated by centrifugation and aldosterone levels were measured using ELISA kit (ENZO, ADI-900-173; sensitivity= 4.7pg/mL; intra- and inter-assay coefficient of variance <6.6% and <18%, respectively).

### Metabolic Cages

To measure the effect of KS-WNK1 on intake, output, and blood and urine parameters, mice were individually housed in metabolic cages (Tecniplast, Italy). Mice were fed pellet-based diets during days 1-7, introduced to powder diets on day 7, and then switched to exclusive powder-diets for days 8-10. Powder diets were derived from blended commercial diets combined with water that was allowed to evaporate to form a solid that was simple to weigh and use in the metabolic cage. The mice were acclimated in metabolic cages for the first 24h (day 9), followed by 24h measurement of food and water intake and urine collection (day 10). Urine [Na^+^], [K^+^], and [Cl^-^] were determined using Easy Lyte Plus Na/K/Cl analyzer (Medica Corp, Bedford, MA). Urine osmolality was determined using a micro osmometer (Precision Systems, Natick, MA). After 10 days mice were anesthetized with isoflurane, and blood was obtained via terminal cardiac puncture and analyzed by iSTAT (Abbot). Kidneys were processed for immunoblot and imaging as stated below.

### Human studies

The Pitt Biospecimen Core provided formalin-fixed, paraffin-embedded non-neoplastic kidney tissue from 6 subjects who underwent radical nephrectomy for renal tumors. An IRB-approved Honest Broker provided the de-identified, safe harbor data including age, sex, and serum K+ measured on the day of surgery. All research was approved by the University of Pittsburgh Internal Review Board (IRB) for secondary research with data and /or specimens.

### Immunohistochemistry

Formalin-fixed, paraffin-embedded human kidney tissue sectioned at 5-μm thickness was processed for immunohistochemistry according to our previously published methods ^12^. Briefly, sections were deparaffinized and hydrated for retrieval of antigenic sites before inactivation of endogenous peroxidases. After blocking, slides were incubated in primary antibody against WNK1 or NCC (see antibodies section) overnight, and subsequent incubation with biotinylated secondary antibodies (Jackson Immunoresearch) and ABC reagent were used to visualize staining with diaminobenzadine (Vector Laboratories). Images were acquired with a Leica DM6000B widefield microscope with a Retiga 4000R Fast 1394 camera. Images were obtained with Volocity 6.3, and analysis was performed using ImageJ/Fiji (NIH).

### Immunoblotting

For protein quantification, kidney cortexes were flash frozen and processed as previously described ^22^. Ice-cold RIPA buffer (Thermo Scientific) was used for protein extraction with freshly added protease and phosphatase inhibitor cocktail. Protein quantification was determined using the Pierce BCA Protein Assay Kit (Thermo Scientific). Uniform protein loading was determined using Coomassie-stained gels, as previously described ^22,48^. 15µg protein from each sample was loaded onto the SDS-PAGE gel and then stained with Coomassie blue. Five random bands were quantified to determine uniform loading, defined as signal variability per lane of less than or equal to 5%. If necessary, protein assays were repeated and appropriate adjustments were made until loading across samples was determined to be uniform. Final Coomassie optimized gels are shown in Figs S1A-F, S2E-G, S7A, and S7B. Next, equal amounts (20-40µg) of protein were separated by SDS-PAGE using 4-20% Criterion TGX precast gels (Bio-Rad). Protein was transferred to a nitrocellulose membrane. Signal densitometry was measured using Bio-Rad ChemiDoc and densitometry was quantified with ImageLab analysis software (Bio-Rad). Two different protein ladders were used: Precision Plus All Blue (Biorad) and PageRuler Plus (Thermo Scientific).

To plot NCC densitometry values as a function of blood [K^+^], WT and mutant mice placed on control diets were run on the same gel as WT and mutant mice treated with a specific potassium maneuver. This permitted normalization of all values to the protein abundance in WT mice on control diet. All normalized densitometry values were disaggregated by sex: males subjected to potassium maneuvers were normalized to WT control diet-treated males, and females subjected to potassium maneuvers were normalized to WT control diet-treated females. The densitometry values for each band – derived from the kidney lysate from one mouse – was cross referenced with the blood potassium concentration measured in that mouse at the time of sacrifice. The normalized densitometry values were then plotted with blood [K^+^] on the X axis and protein abundance on the Y axis.

### Antibodies

The following antibodies were used for immunoblot and/or immunofluorescence: Sodium chloride cotransporter (NCC; provided by David Ellison^49^); Sodium chloride cotransporter (NCC; Millipore Ab3553^50^); Phospho-NCC Thr 53 (pNCC; Phospho-solutions P1311-53^51^); Phospho-NCC Ser 71 and Phospho-NCC Thr 58 (pNCCSer71 & pNCCThr58, provided by Jan Loffing^4^); SPAK (SPAK; Cell Signaling^50^); Phospho-Ser 373 SPAK/ Phospho-Ser 325 OSR1 (pSPAK/pOSR1; Millipore 07-2273^14^); With-no-lysine kinase 1 (WNK1; Atlas Antibodies HPA059157^12^); With-no-lysine kinase 4, (WNK4^12^); Gamma subunit epithelial sodium channel (γENaC; 83 kDa cleaved and 95 kDa uncleaved; Stressmarq SPC-405)^22^; ATP-sensitive inward rectifier potassium channel 10 (Kir4.1; Alomone APC-035^52^); Renal outer medullary potassium channel (ROMK; 50-65 kDa complex glycosylated; R-80 was provided by James Wade^53^).

### Blood Pressure Telemetry

Female KS-WNK1-KO mice and WT littermates were anesthetized with isoflurane and DSI PA-C10 telemetry units (Data Sciences International, New Brighton, MN, USA) were surgically implanted into the femoral artery as previously described^54^. Mice then recovered for 1 week before dietary challenges were commenced. Blood pressure was collected every day from 10am-4pm (daytime) and 10pm-4am (nighttime) for the duration of the diet challenge using Spike2 software (Cambridge Electronic Design). Mean arterial pressure (MAP) was calculated by diastolic plus one-third of the pulse pressure. For the saline challenge, mice were maintained on varying K+ diets for 10 days and were then challenged with 1% saline in their drinking water for 72h. HCTZ-challenge was performed 72h after saline administration commenced. During the HCTZ treatment mice were maintained on low K+ diet with 1% saline drinking water. Daytime blood pressure was obtained for 6h on 3 separate days: 1) the day prior to HCTZ administration, 2) the day of HCTZ administration, and 3) the day after HCTZ administration. Mice were injected with hydrochlorothiazide (25mg/kg IP) at 9am and then blood pressure was collected from 10am-4pm (daytime).

### Diuretic Challenge

Mice were placed on respective K^+^ diets for 10 days. For the first 8 days they were housed in the standard cages. On day 9 they were individually housed in metabolic cages and acclimated for 24h. On day 10 mice were given intraperitoneal injections of hydrochlorothiazide (HCTZ; 25mg/kg) and urine was collected for 6 hours. Urine [Na^+^], [K^+^], and [Cl^-^] were determined using Easy Lyte Plus Na/K/Cl analyzer (Medica Corp, Bedford, MA).

### Quantitative Immunofluorescence

Paraformaldehyde fixed kidney tissues were processed and prepared as previously described^22^. 6μm sections were rehydrated and treated with 1% SDS for 10 min for retrieval of antigenic sites. Then slides were washed with high salt buffer + bovine serum albumin before the addition of primary antibody. Primary antibodies were incubated overnight at 4°C, followed by washes with high salt buffer + bovine serum albumin and subsequent incubation with secondary antibodies and TO-PRO-3 Iodide for visualize staining. Imaging of the kidney tissue was performed using a Leica HCX PL APO CS x40, 1.25 numerical aperture oil objective on a Leica TCS SP5 CW-STED confocal microscope utilizing Leica LAS-X software.

To produce quantitative measures for pSPAK/pOSR1 puncta number, size, and distance to the DCT lumen we used Imaris (Bitplane, v9.5) image analysis software. Fluorescence images were first imported into Imaris. The pSPAK/pOSR1 puncta were detected using the Spots element creation wizard. A region of interest was specified to isolate a single DCT within an image. The pSPAK/pOSR1 channel was then selected as the fluorescence reference channel for both Spot identification and to guide the puncta diameter determination. A local contrast fluorescence intensity threshold was chosen, and the Spots elements were filtered using the Quality metric intrinsic to Imaris. The region growing method was used to obtain Spots of varying size. Once defined, the same quality metric and threshold settings were used throughout all puncta analysis. To not exclude data, the quality and thresholding was set to bias towards over detection. Manual editing of the Spot objects was performed to eliminate rare errant Spots when necessary. For each DCT the number of cell nuclei were counted and used to calculate the average number of pSPAK/pOSR1 puncta per cell. To measure the distance between the identified spots and the DCT lumen, we created a Surface object of the lumen. Using the Surface wizard, a Surface object was defined using the magic wand and isolines functions within the manual surface creation. This was able to reliably detect the lack of fluorescence intensity within the DCT lumen. A built-in Imaris Xtension was implemented to calculate the shortest distance between all Spots objects and the DCT lumen Surface. These data were then displayed in Imaris as Spot objects with color coded statistic values for puncta diameter or distance to DCT lumen. To measure WNK condensate morphology in 5Q mice, confocal images of WNK1 signal from WT and KS-WNK1 5Q mice were obtained under identical confocal settings. Images were thresholded under identical parameters to generate masks (Fig 7B), which were then used to measure the area and roundness of individual puncta, using the “Analyze Particles” tool in FIJI.

For WNK body analysis and quantification in formalin-fixed, paraffin-embedded mouse kidney, tissues sectioned at 5-μm thickness were processed for immunofluorescence staining according to a protocol similar to the immunohistochemistry protocol described above. After a 1-hour blocking step with donkey serum, slides were incubated in primary antibodies against WNK4 or NCC (see antibodies section) overnight, and a subsequent 2-hour incubation with fluorescent-tagged secondary antibodies (Jackson Immunoresearch) was used to visualize staining. Images were acquired with a Leica DM6000B widefield microscope with a Retiga 4000R Fast 1394 camera. Up to ten images of representative distal convoluted tubules (DCTs) per animal were captured using ImageJ. Two independent, blinded analyses were conducted to measure WNK body size and count using a custom macro in ImageJ. Threshold values were adjusted to exclude background staining, with the same settings applied consistently across tubules from the same animal. The results from both analyses were averaged, unblinded, and plotted as a function of blood [K^+^].

### Protein structure prediction

The amino-terminal structures of WT and 5Q mutant KS-WNK1 (residues 1-72, encompassing exon 4a to the C-terminal end of the remnant kinase domain; residue 494 of Uniprot sequence Q9JIH7-1) were predicted using ColabFold^55^, accessed via UCSF ChimeraX^56^. Full-length L- and KS-WNK1 were rendered with AlphaFold3^57^.

### Data analysis

Data were analyzed using GraphPad Prism software and presented as mean ± SE, plus individual data points. Comparisons between two groups were determined by a Student’s T-test. Comparisons between multiple groups were determined using one- or two-way analysis of variance (ANOVA), followed by the appropriate post hoc test, as indicated. P values ≤-s 0.05 were considered statistically significant. To analyze the relationship between blood [K^+^] and NCC, regression analyses were performed in Prism. Data were preliminarily fitted via one phase decay, indicating an exponential relationship with suboptimal goodness of fit. Thus, the absolute Y-values (representing normalized protein densitometry as described above) were logarithmically transformed and analyzed by segmental vs simple linear regression. A comparison of fits was performed by F-test, with data presented fit to the preferred regression model (alternative hypothesis = segmental vs null hypothesis = straight line). P values for these comparisons are shown with the corresponding residuals in Fig S4.

## Supplemental Information

### SI Table of Contents

#### I. Supplemental Figures

**Figure S1.**
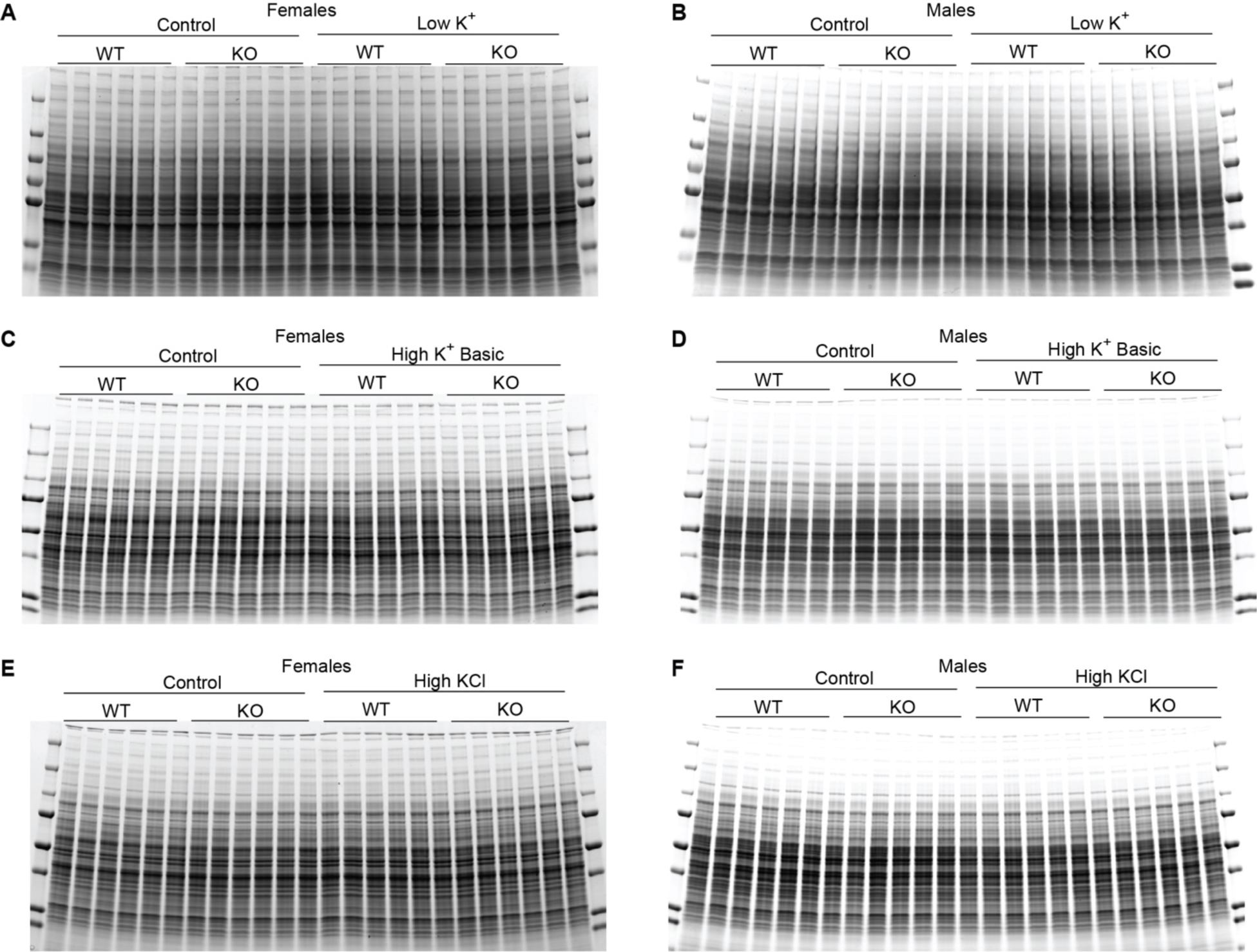
Optimized Coomassie gels for immunoblotting (KS-WNK1 KO experiments). Coomassie blue stained gels, optimized to verify equal protein loading. To make comparisons between different diets, immunoblots were run against the same lysate from control WT samples in the first 12 lanes. Gels correspond to immunoblots presented in multiple figures, as noted above. **A-F.** (A) Female WT and KS-WNK1 KO mice on control vs low K^+^ diet. (B) Male WT and KS-WNK1 KO mice on control vs low K^+^ diet. (C) Female WT and KS-WNK1 KO mice on control vs HKB diet. (D) Male WT and KS-WNK1 KO mice on control vs HKB diet. (E) Female WT and KS-WNK1 KO mice on control vs HKCl diet (F) Male WT and KS-WNK1 KO mice on control vs HKCl diet.

**Figure S2.**
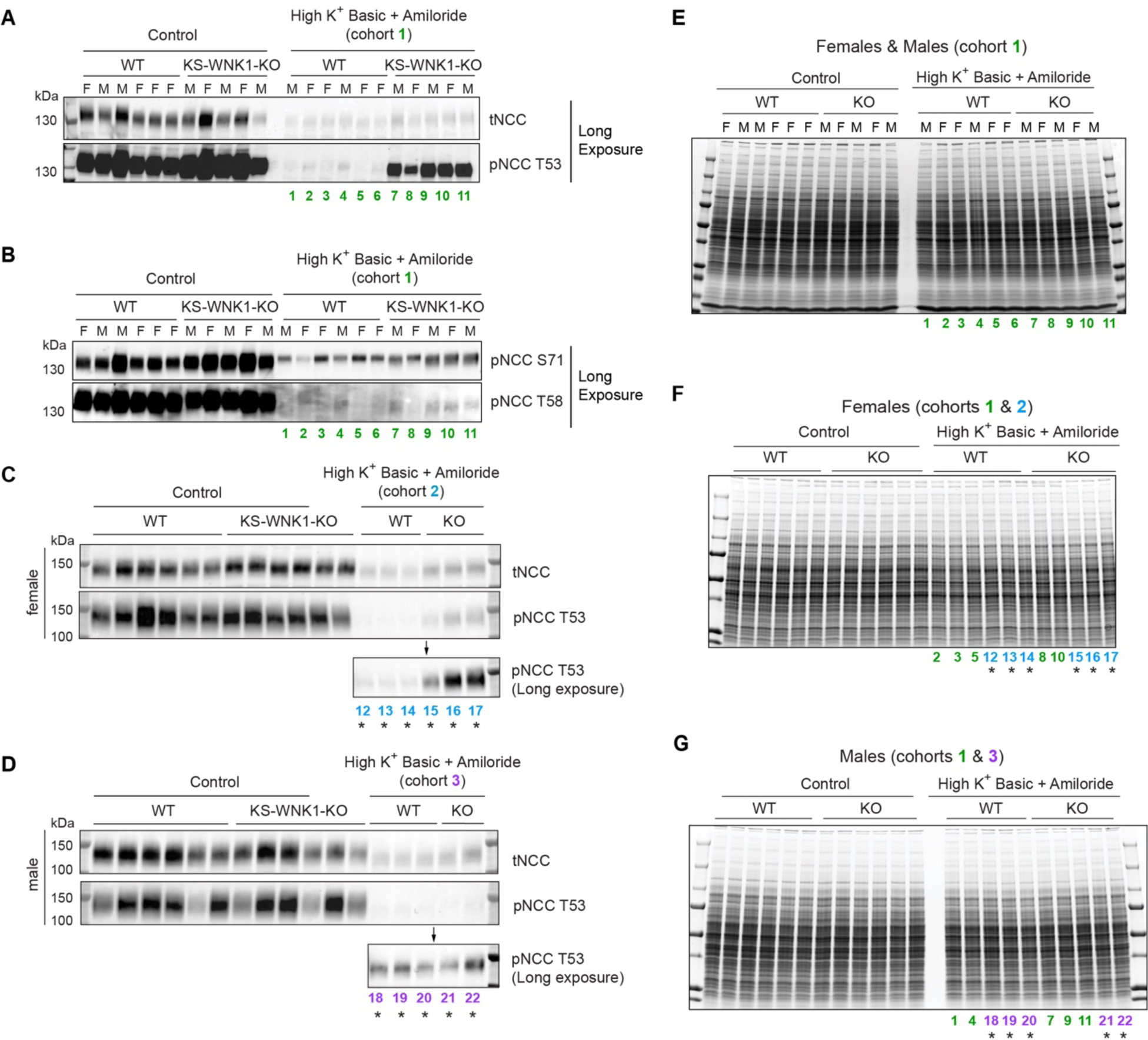
Effect of potassium loading + amiloride on NCC phosphorylation in KS-WNK1 KO mice. Immunoblotting data for mice subjected to alkaline K^+^ loading with amiloride (2mg/kg/day) were derived from 3 separate cohorts. Cohort 1 (male + female) was studied first. Cohorts 2 (female only) and 3 (male only) were studied several months later. These additional cohorts were included to achieve sufficient *n* for disaggregation of results by sex. *n* was limited for all cohorts due to extreme hyperkalemia (Table S2). Mice from cohorts 1, 2, and 3 are indicated in green, blue, and purple, respectively. Lysates from individual mice were numbered as indicated (*i.e.,* #1 represents the same mouse lysate in all blots). The increased signal in the KO mice treated with the HKB diet + amiloride was conspicuous after prolonged membrane exposure, as shown. **A.** Cohort 1, with male and female protein lysates analyzed on the same blot. Male (M) mice were normalized to male controls, and female (F) mice were normalized to female controls. Phosphorylated NCC was probed with pNCC-Thr53 antibody. **B.** pNCC-Thr58, and pNCC-Ser71 antibodies also detected an increase in pNCC in KS-WNK1 KO mice subjected to HKB + amiloride. **C.** Cohort 2, female mice treated with HKB + amiloride. **D.** Cohort 3, male mice treated with HKB + amiloride. **E-G.** Optimized Coomassie gels. Lanes indicated by asterisks (*) correspond to the mice that were used for the sex-disaggregated tNCC and pNCC immunoblots in C & D.

**Figure S3.**
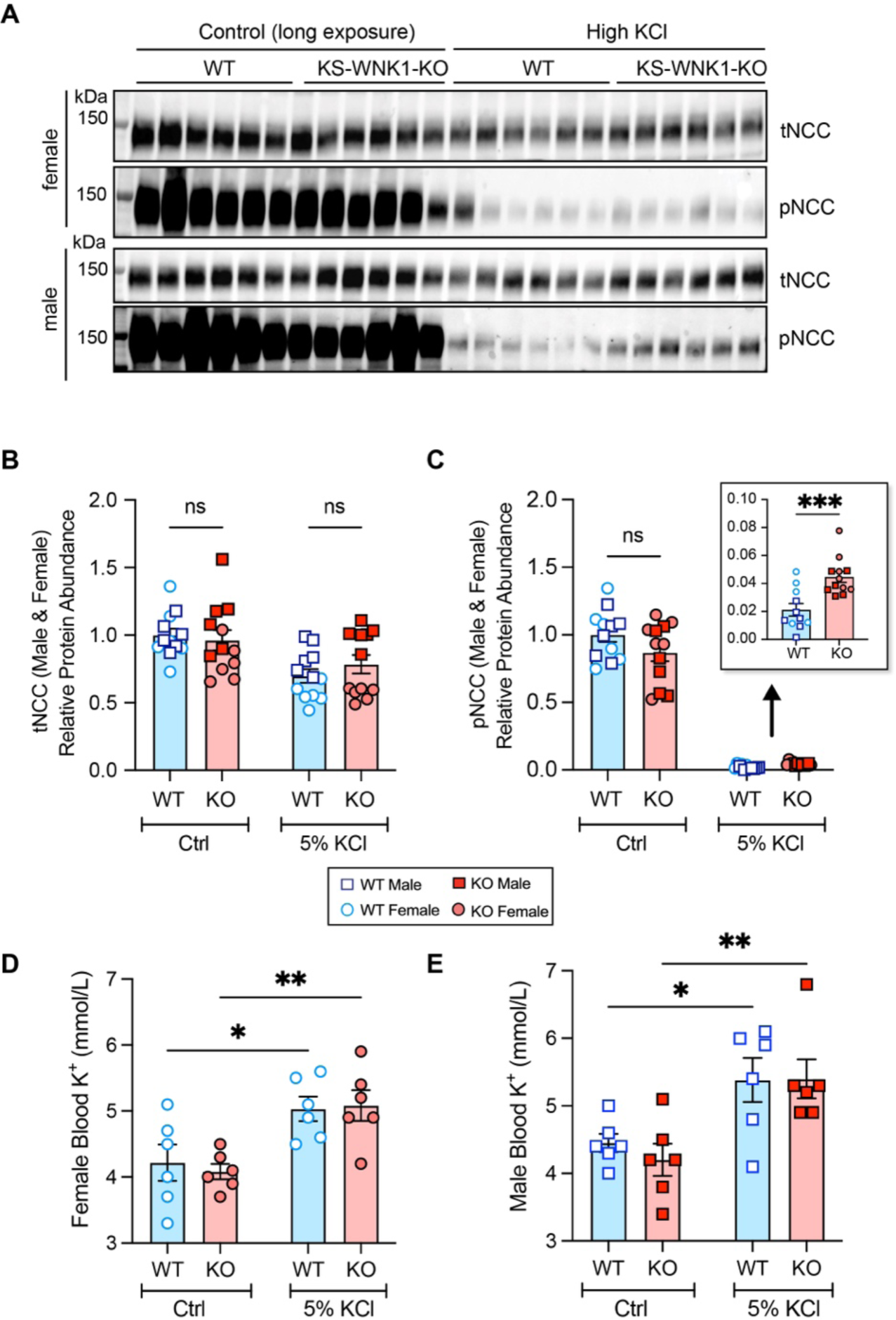
Effect of high KCl diet and KS-WNK1 on NCC activation. **A.** Immunoblot of kidney cortical extracts from mice treated with either control or high KCl diet for 10 days. Control diet results are shown overexposed to visualize high KCl results. **B-C.** Quantification of (A). High KCl diet significantly reduced pNCC abundance. KS-WNK1 KO mice have a blunted reduction in pNCC abundance compared to WT mice on high KCl diets. **D-E.** Whole blood [K^+^] in mice treated with high KCl diets. Results are shown as means ± SE; *n* = 5-6 mice per genotype per diet. Two-way ANOVA with Sidak’s multiple comparisons test was applied comparing WT and KS-WNK1 KO, \**P* ≤ 0.05, \*\**P* ≤ 0.01, \*\*\**P* ≤ 0.001, \*\*\*\**P* ≤ 0.0001.

**Figure S4.**
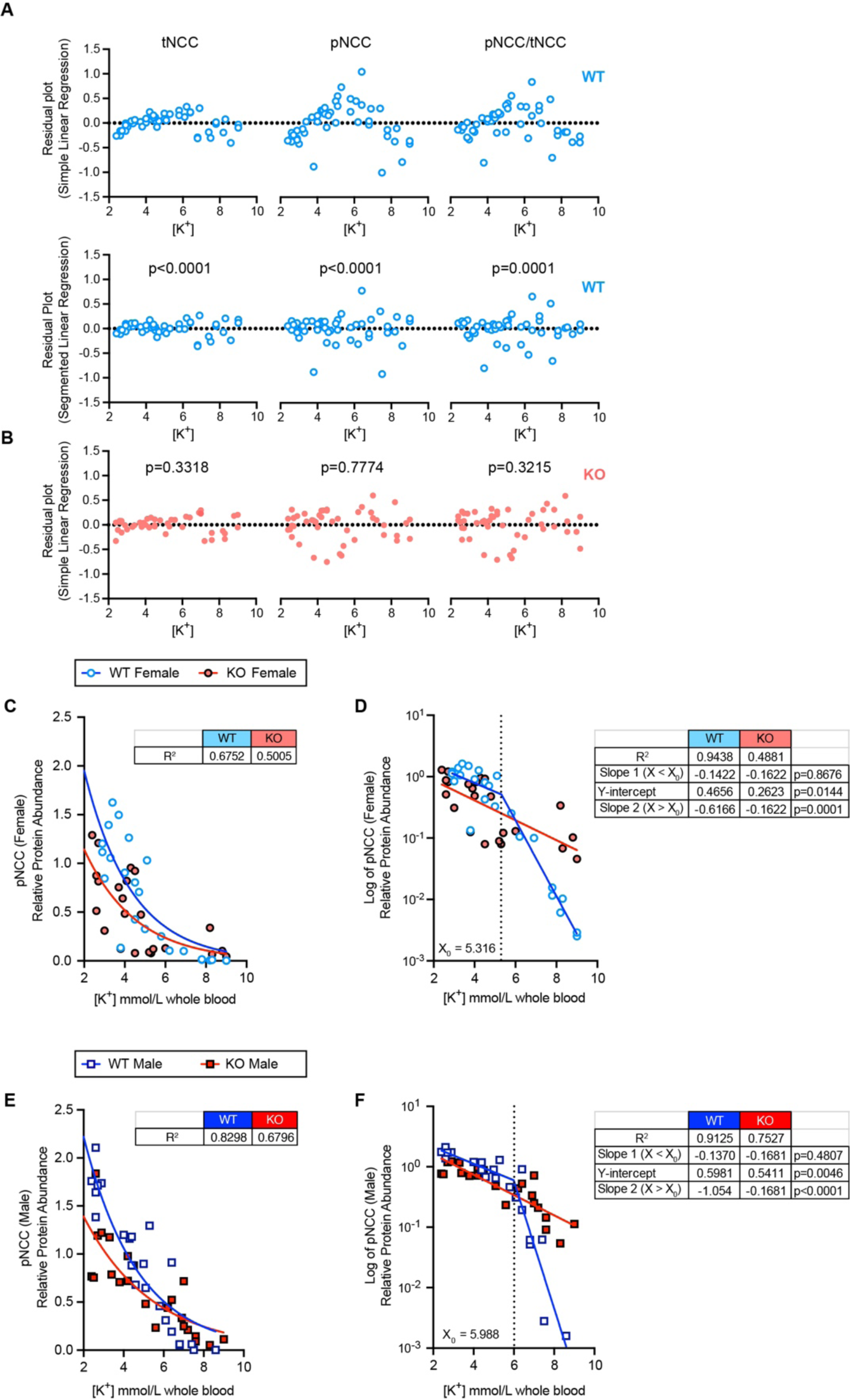
Residual plots for log transformed NCC data versus [K+] in WT and KS-WNK1 KO mice. **A.** Residual vs X plots for log-transformed tNCC, pNCC, and pNCC/tNCC data in WT mice, modeled by simple linear regression (top) or by segmented linear regression (bottom). For the simple linear regression residuals, note how all three plots exhibit a nonrandom concave-down distribution relative to the horizontal dotted line, suggesting systematic inconsistencies in the models and, therefore, suboptimal goodness-of-fit. When the data were re-modeled by segmental linear regression, the residuals for all three datasets were distributed normally. P-values indicate the results of a comparison-of-fits between the segmented vs simple linear models, with simple linear regression as the null hypothesis (P ≤ 0.0001 for all three WT datasets). **B.** Residual vs X plots for log-transformed tNCC, pNCC, and pNCC/tNCC data in KS-WNK1 KO mice, modeled by simple linear regression. P-values indicate the result of a comparison-of-fits test between segmented vs simple linear regression models, with simple linear regression as the null hypothesis. In the case of KS-WNK1 KO mice, segmental linear regression did not improve the fit for KO animals and was therefore rejected, as indicated by the non-significant P-values. **C,E.** pNCC in female and male mice, fit to single exponential curves. R^2^ measures are presented in table format alongside the graphs. **D, F.** Normalized pNCC densitometry in A & B were log transformed and analyzed by linear regression. WT data were bet fit by a segmented linear regression regime, with X0 breakpoints (dotted line) shown. Slopes of the two linear components are presented in table format alongside the corresponding graphs. Consistent with the combined analysis of male and female data, For KO mice, the log-transformed data were best fit by simple linear regression. P-values represent slope comparisons between WT and KO; in the event where Slope 1 (X < X0) comparisons did not reach significance, Y-intercept comparisons with P-values are shown.

**Figure S5.**
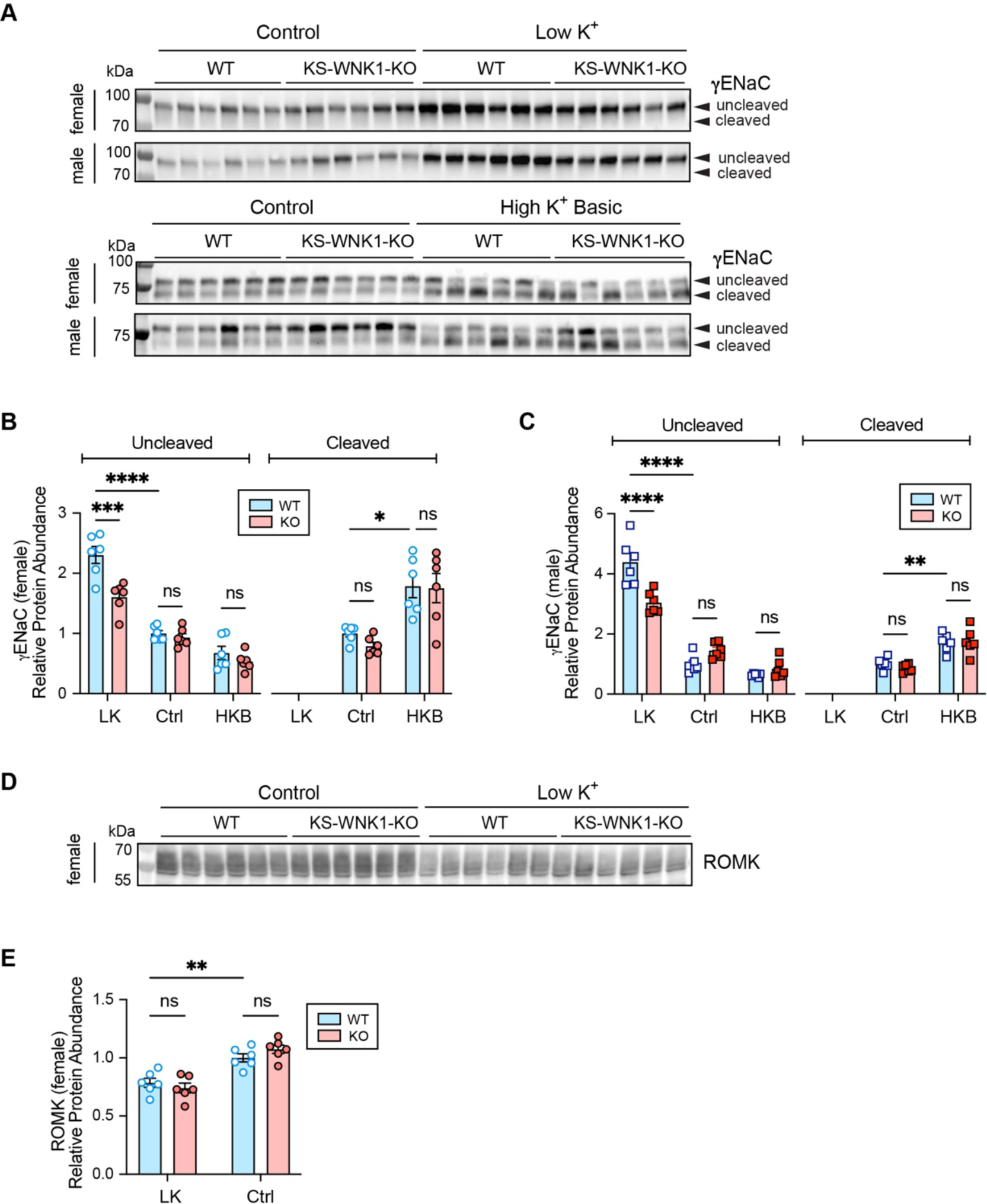
Effects of KS-WNK1 and dietary potassium on ENaC and ROMK protein abundance. IB analysis of kidney cortical extracts from female and male WT and KS-WNK1 KO mice fed low K+, control, and high K+ basic for 10 days. **A.** γENaC. Both the cleaved (hyperactive)^1^ and uncleaved forms are indicated with an arrow. **B & C.** Bar graphs showing relative protein abundance, normalized to WT littermates on control diet. The KO mice have significantly reduced uncleaved γENaC during low K^+^ diet. The cleaved form of γENaC is undetectable during low K^+^ diet. **D & E.** Low K^+^ diet reduces the abundance of ROMK in female mice, however KS-WNK1 expression has no effect on ROMK protein abundance. Results are shown as means ± SE; *n* = 5-6 mice per genotype per diet. Two-way ANOVA with Sidak’s multiple comparisons test was applied comparing WT and KS-WNK1-KO, \**P* ≤ 0.05, \*\**P* ≤ 0.01, \*\*\**P* ≤ 0.001, \*\*\*\**P* ≤ 0.0001.

**Figure S6.**
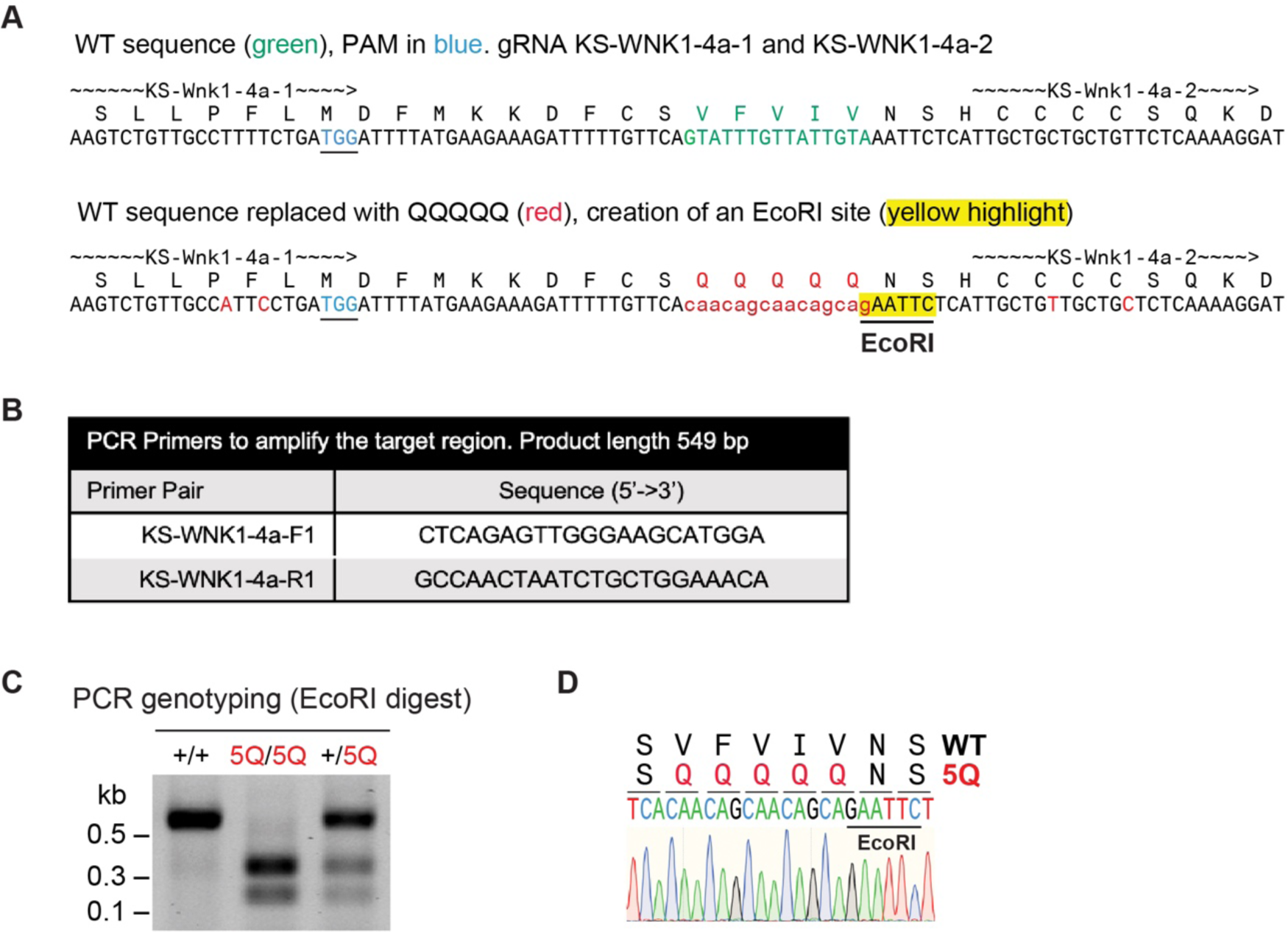
Generation of KS-WNK1 5Q mutant mice. **A.** The Exon 4a encoded bulky hydrophobic patch VFVIV of the CRH motif (shown in green) was replaced with QQQQQ (shown in red) via homology-directed repair (HDR). The HDR template also contained an engineered EcoRI site (yellow highlight) for genotyping purposes. Single guide RNAs flanking the hydrophobic domain are indicated as KS-WNK1-4a-1 and KS-WNK1-4a-2. The guides were injected into 129-Elite Mouse 129S2/SvPasCrl fertilized embryos, which were transferred to pseudo-pregnant female recipients. **B.** Pups were then genotyped by PCR using the indicated primers. **C.** PCR gel with genotyping via EcoRI digest indicating homozygous 5Q/5Q mutant mice. The founders were backcrossed with 129-Elite Mouse 129S2/SvPasCrl for 5 generations prior to experimentation. **D.** Sanger sequencing of the TA-cloned genomic DNA PCR product confirming the correct mutation with diagnostic EcoRI site in 5Q mutant mice.

**Figure S7.**
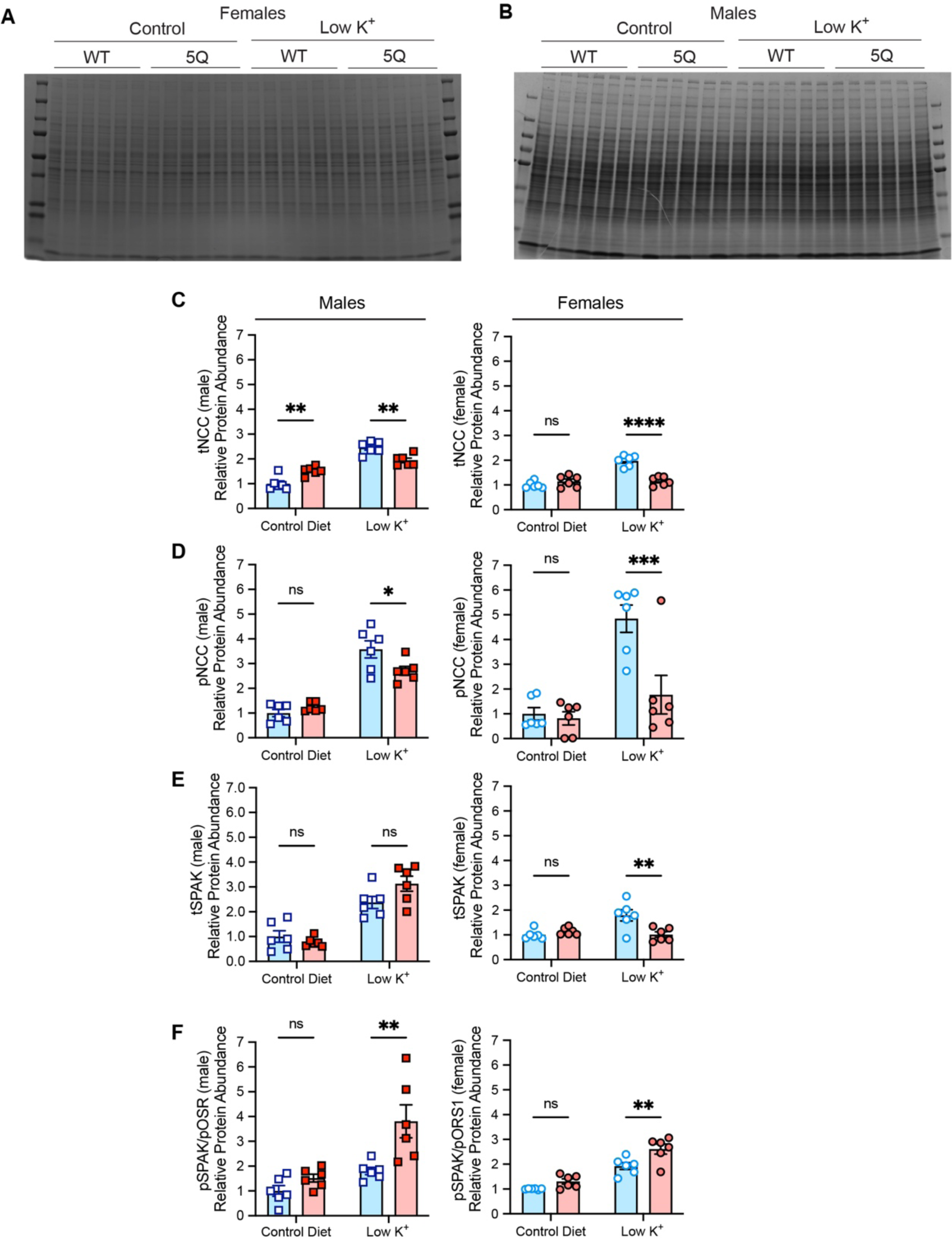
Potassium-restricted KS-WNK1 5Q mice: optimized Coomassie gels and data quantification disaggregated by sex. **A-B.** Coomassie blue stained gels, optimized to verify equal protein loading; (A) Female WT and KS-WNK1 5Q mice on control vs low K^+^ diet, (B) Male WT and KS-WNK1 5Q mice on control vs low K^+^ diet. **C-F.** tNCC, pNCC, tSPAK, and pSPAK/pOSR1 protein abundance in separated female and male KS-WNK1 5Q (red) versus WT (blue) littermates. (C) tNCC abundance in female and male mice (D) pNCC abundance in female and male mice. (E) tSPAK abundance in female and male mice. (F) pSPAK/pOSR1 abundance in female and male mice. Two-way ANOVA with Sidak’s multiple comparisons test was applied comparing WT and KS-WNK1 5Q mutants, \**P* ≤ 0.05, \*\**P* ≤ 0.01, \*\*\**P* ≤ 0.001, \*\*\*\**P* ≤ 0.0001.

#### II. Supplemental Tables

**Table S1.** Mineral content in the various K^+^ diets used in the study.

| Table S1: Calculated percent of minerals in the varying K<sup>+</sup> diets |  |  |  |  |
| --- | --- | --- | --- | --- |
|  | Low K<sup>+</sup><br>TD.88238 | Control<br>TD.88239 | High K<sup>+</sup> Basic<br>TD.07278 | High KCl<br>TD.88239 |
| Na<sup>+</sup> | 0.3% | 0.3% | 0.3% | 0.3% |
| K<sup>+</sup> | 0.003% | 1% | 5%* | 5% |
| Cl<sup>-</sup> | 0.45% | 0.9% | 2% | 5.2% |
| Ca<sup>2+</sup> | 1% | 1% | 1% | 1% |
| *with a 1:1:1 ratio of K<sup>+</sup> citrate, KCl, K<sup>+</sup> carbonate |  |  |  |  |

**Table S2.**
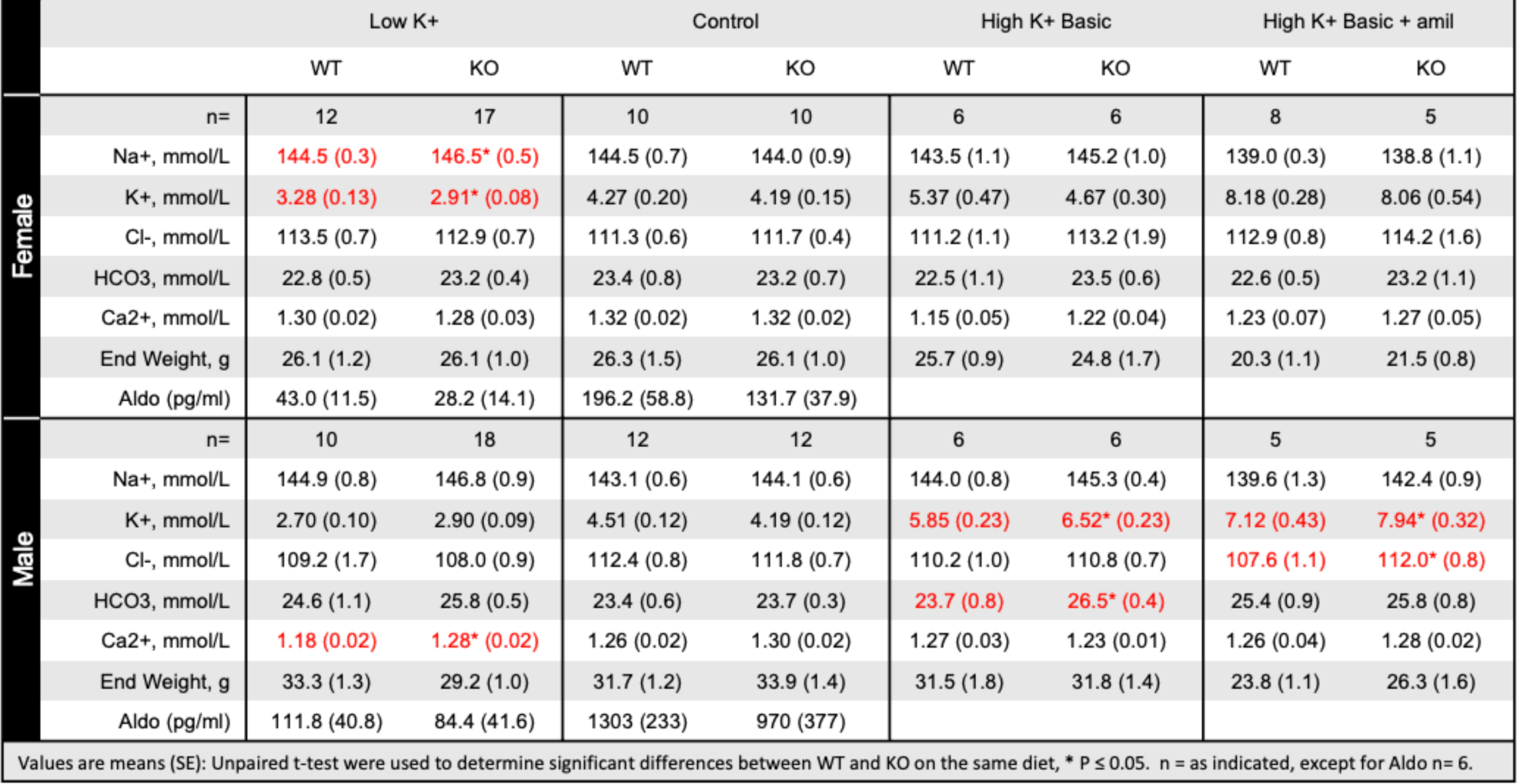
Effects of KS-WNK1 deletion on whole blood parameters in mice on varying K^+^ diets.

**Table S3.**
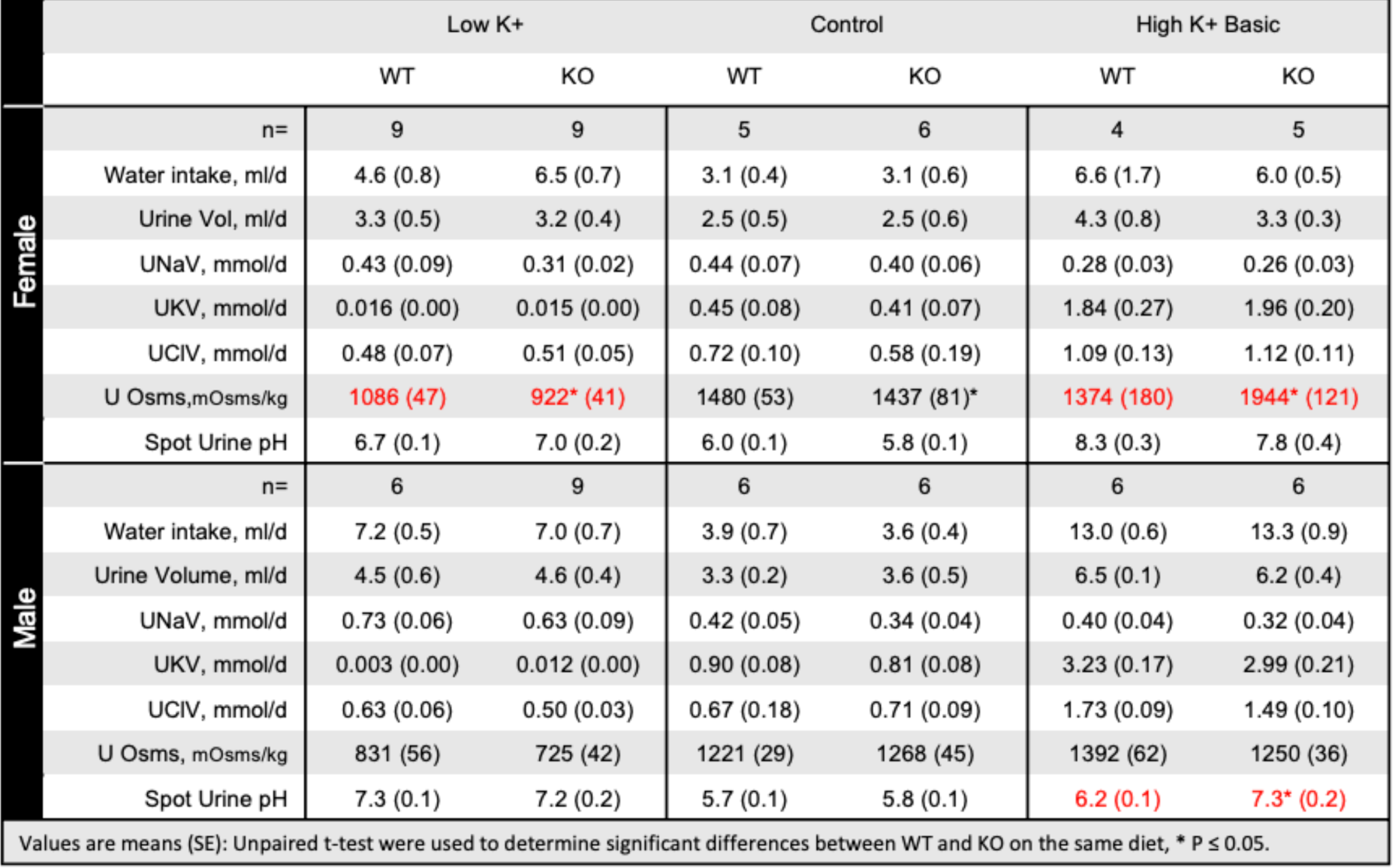
Effects of KS-WNK1 deletion on urine parameters in mice on varying K^+^ diets.

**Table S4.**
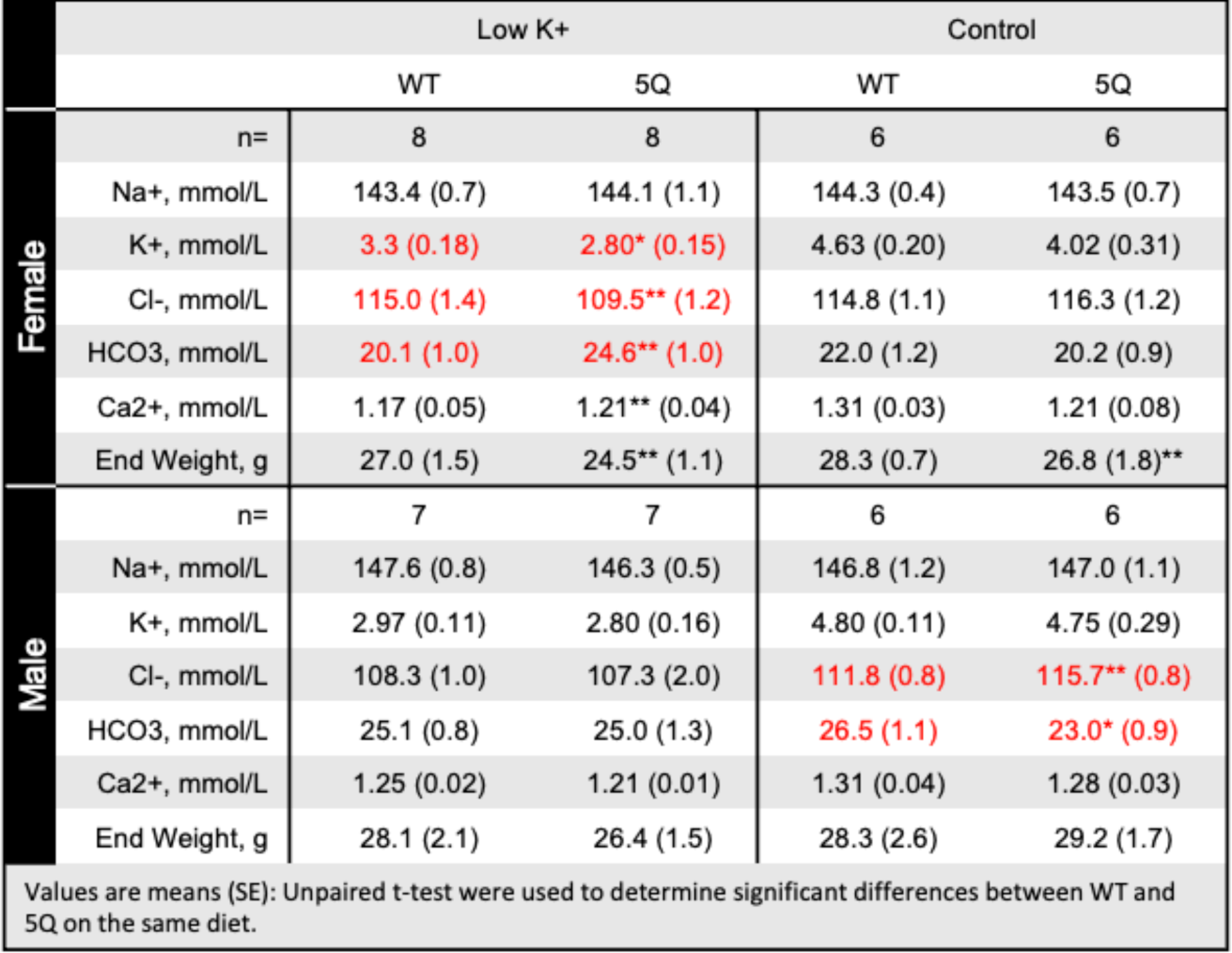
Effects of the KS-WNK1 5Q mutation on whole blood parameters in mice on varying K^+^ diets.

